# Stable and ancient endocytic structures navigate the complex pellicle of apicomplexan parasites

**DOI:** 10.1101/2022.06.02.494549

**Authors:** Ludek Koreny, Brandon N. Mercado-Saavedra, Christen M. Klinger, Konstantin Barylyuk, Simon Butterworth, Jennifer Hirst, Yolanda Rivera-Cuevas, Nathan R. Zaccai, Victoria J. C. Holzer, Andreas Klingl, Joel B. Dacks, Vern B. Carruthers, Margaret S. Robinson, Simon Gras, Ross F. Waller

## Abstract

Apicomplexan parasites have an immense impact on humanity, but their basic cellular processes are often poorly understood. The sites of endocytosis, the conservation of this process with other eukaryotes, and its functions across Apicomplexa are major unanswered questions. Yet endocytosis in *Plasmodium* is implicated in antimalarial drug failure. Using the apicomplexan model *Toxoplasma*, we identified the molecular composition and behavior of unusual, fixed endocytic structures. Here, stable complexes of endocytic proteins differ markedly from the dynamic assembly/disassembly of these machineries in other eukaryotes. Moreover, conserved molecular adaptation of this structure is seen in Apicomplexa, including the kelch-domain protein K13 central to malarial drug-resistance. We determine that an essential function of endocytosis in *Toxoplasma* is plasma membrane homeostasis, rather than parasite nutrition, and that these specialized endocytic structures originated early in infrakingdom Alveolata, likely in response to the complex cell pellicle that defines this medically and ecologically important ancient eukaryotic lineage.

## INTRODUCTION

Apicomplexa is a diverse group of eukaryotic intracellular parasites infecting every major animal taxon including humans. The malaria-causing *Plasmodium* spp. are responsible for over 600,000 deaths a year (World Health Organization, 2021), *Cryptosporidium* spp. are a leading cause of diarrhea morbidity and mortality in children younger than 5 years (Khalil et al., 2018; Striepen, 2013), and *Toxoplasma gondii* is the most prevalent human parasite estimated to infect a third of the world’s population (Havelaar et al., 2015). While in most adults *Toxoplasma* does not cause serious illness, it can cause life-threatening congenital toxoplasmosis, fetal malformation and abortion, blindness, and encephalitis, with immunocompromised individuals most susceptible (Daher et al., 2021; Montoya and Liesenfeld, 2004). Other members of Apicomplexa infect economically important livestock through which they also have major impact on human wellbeing (MacGregor et al., 2021).

A key feature of apicomplexans is the inner membrane complex (IMC) — a near continuous array of flattened membranous vesicles (alveoli) underneath the plasma membrane and supported by a complex proteinaceous membrane skeleton. This IMC pellicle provides cell shape and strength, and is the critical platform for a gliding motility apparatus that allows parasite tissue traversal and the mechanism for host cell invasion (Frénal et al., 2017; Harding and Meissner, 2014). The IMC, however, separates most of the parasite’s plasma membrane from its cytoplasm and is, therefore, a barrier to the material exchange processes of endocytosis and exocytosis. This is an ancient problem for these cells because the alveolae-based pellicle predates the development of apicomplexan parasitism as it is a common feature shared with dinoflagellates and ciliates, the other two major lineages of the infrakingdom Alveolata (Gould et al., 2008; Kono et al., 2013). Understanding the solutions to the challenges presented by the IMC are, therefore, equally important to understanding the major ecological functions of these organisms in ocean primary production, coral symbiosis, food webs and nutrient recycling. Adaptations for exocytosis across the IMC are best studied in Apicomplexa because regulated discharge of secretory organelles micronemes and rhoptries drive the processes of host cell attachment, motile exploration, and invasion of their host cells (Dubois and Soldati-Favre, 2019; Joiner and Roos, 2002; Sparvoli and Lebrun, 2021). This secretion occurs through the apical complex, a cytoskeletal feature integrated into the IMC that provides a window of available plasma membrane at the cell apex for vesicle docking and fusion (dos Santos Pacheco et al., 2020). Considerable insight into the molecular details of these exocytic processes has been achieved through decades of intense investigation (Kremer et al., 2013; Mageswaran et al., 2021; Sloves et al., 2012), and it is evident that many of these adaptations are present in the related dinoflagellates and ciliates (Koreny et al., 2021). The details and adaptations for endocytosis, on the other hand, have received far less attention in these organisms.

The importance of endocytosis in apicomplexans has been brought into sharp focus in recent years through a link to the emergence and spread of *Plasmodium* tolerance to the lead antimalarial drug artemisinin and its derivatives (ARTs). Many ART-resistant *Plasmodium* field isolates have correlated with mutations in either a kelch (beta-propeller) domain of a protein called Kelch13 (K13) or the medium (μ) subunit of the AP-2 adaptor complex (AP-2μ) (Henriques et al., 2015; Miotto et al., 2015). In the mouse model for malaria, *P. berghei*, ART-resistant parasites selected in the laboratory similarly contain mutations in AP-2μ, as well as in a deubiquitinase UBP1 (Henriques et al., 2013). While the functional significance of these mutations was initially unclear K13, AP-2μ and UBP1, along with 11 other proteins, were recently shown to associate with the cytostome of the feeding blood-stage of *Plasmodium* (Birnbaum et al., 2020; Xie et al., 2020; Yang et al., 2019b). The cytostome is a prominent membrane invagination where hemoglobin-rich host cytoplasm is taken up for digestion. Loss of function of five of these proteins at the cytostome were all seen to correlate with reduction in hemoglobin uptake, and the inactivation of eight of them conferred increased ART-tolerance. The hemoglobin degradation pathway is important for the activation of ARTs, and decreased levels of ingested hemoglobin in the mutant strains is believed to be main cause of the resistance (Birnbaum et al., 2020; Yang et al., 2019b). The cytostome, however, is specific to the feeding-stages of erythrocytic *Plasmodium* where the IMC is temporarily disassembled. Furthermore, of the fourteen identified K13 complex proteins in *Plasmodium* (Birnbaum et al., 2020), only AP-2μ and Eps15 are related to previously characterized endocytic factors of other eukaryotes, although the other three canonical AP-2 subunits were enriched in AP-2μ pull-downs consistent with the tetrameric AP-2 adaptor being part of the complex (Henrici et al., 2020). The sites, machinery, or functions of endocytosis in other *Plasmodium* stages, or other apicomplexans, is largely unknown including whether they share any features of the cytostome. Small membrane invaginations named micropores have been observed throughout Alveolata by electron microscopy (Appleton and Vickerman, 1996; Mylnikov, 2009; Nichols et al., 1994), but their hypothesised role in endocytosis is untested and the proteins associated with the micropore are unknown.

In *Toxoplasma*, vesicular uptake from the parasite’s environment has been recorded either as recycling of the surface molecule SAG1 in motile extracellular-stage tachyzoites, or uptake of proteins from the host cell cytoplasm to a lysosome-related vacuole in intracellular tachyzoites (Gras et al., 2019; McGovern et al., 2018). The sites or mechanism of uptake at the parasite plasma membrane, however, are not known and neither are the consequences of the abatement of these processes. Indeed, the significance of endocytosis in *Toxoplasma* is as poorly understood as it is throughout Apicomplexa. In this study we have asked where endocytosis occurs in *Toxoplasma*, what is its major function in this parasite, and does it share mechanistic features with the *Plasmodium* cytostome. We have defined a remarkably stable endocytic complex with molecular conservation to that of the *Plasmodium* cytostome as well as additional endocytic factors previously unknown in Apicomplexa. This complex occurs at two to three sites integrated into the IMC that are preassembled in the IMC well before their endocytic functions are needed. We have developed a new endocytosis assay in *Toxoplasma* that confirms endocytic activity at these sites, and we observe that a major role for this process is maintenance of plasma membrane homeostasis. Moreover, this complex shows conserved molecular adaptations unique to Alveolata implying that it too represents an ancient adaptation that has contributed to the success of this medically and environmentally important lineage of eukaryotes.

## RESULTS

### A K13-associated endocytic complex is present in *Toxoplasma* tachyzoites

Both K13 and the canonical subunits of the AP-2 adaptor complex have orthologues throughout Apicomplexa, but it was unknown if a molecular structure equivalent to the *Plasmodium* cytostome occurs in other apicomplexans. To test for such a structure, we determined the location and associations of these proteins associated with *Plasmodium* ART-resistance in *T. gondii* tachyzoites. We observed reporter-tagged *Tg*K13 (TGME49_262150) located at the periphery of the cell forming multiple (typically 2-3) discrete rings in the plane of the cell surface (Fig 1, S1). The AP-2 adaptor complex is typically comprised of four protein subunits (α, β, μ and σ) where β, in most eukaryotes, is shared with the AP1 adaptor complex which regulates polarized sorting at the trans-Golgi network (Venugopal et al., 2017). While *T*. g*ondii* has single orthologues for the β, μ and σ subunits (TGME49_240870, TGME49_230920, TGME49_313450, respectively), we found two unusually divergent proteins that shared similarity to the canonical α subunit. One protein (designated here *Tg*AP-2α, TGME49_221940) shares similar size and strong sequence identity to the canonical AP-2α proteins but has an unusually divergent C-terminal ‘ear domain’ that typically mediates cognate AP-2 interactions with other proteins. The second *T. gondii* protein contains the highly conserved C-terminal ear domain but the rest of the protein lacks similarity to AP-2α. We designate the second protein KAE (<u>K</u>13 complex-associated, <u>a</u>lpha-<u>e</u>ar containing protein, TGME49_272600). All five of these AP-2-related proteins co-locate with *Tg*K13, with the β subunit displaying additional location within the cell consistent with a shared role with AP-1 (Fig 1A-E, S1, S2A) (Venugopal et al. 2017). The AP-2 components locate within the lumen of the K13 ring presenting as a tapered funnel in cross-section by 3D-SIM super-resolution microscopy (Fig S2A). These data are consistent with endocytic pits lined by AP-2 at the *T. gondii* plasma membrane within a K13-associated ring.

**Figure 1.**
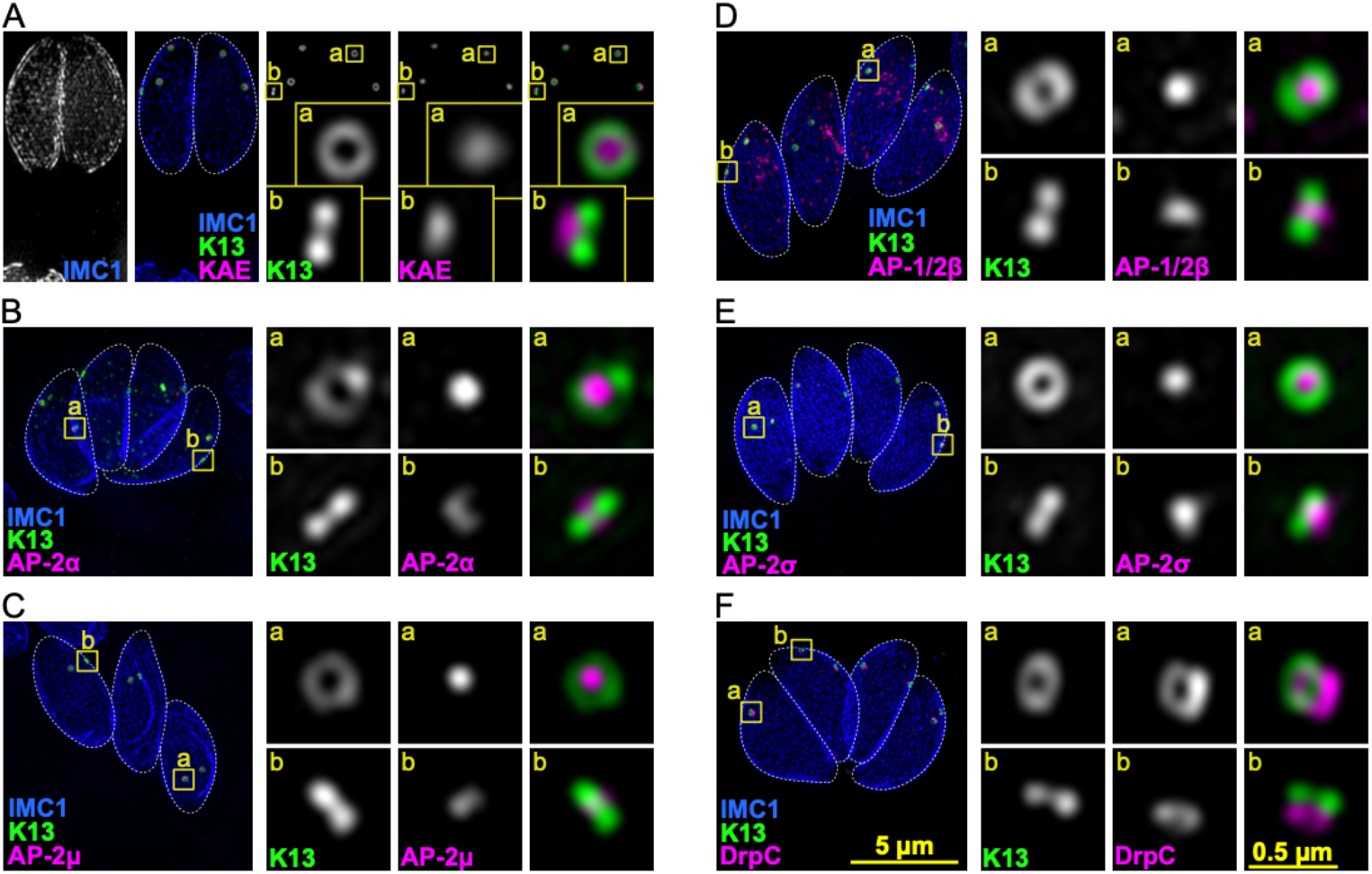
Endocytosis-related AP-2 and DrpC associate with K13 at peripheral circular structures in *T. gondii*. ***A***. Collapsed projections of 3D-SIM images of intracellular parasites within the host cell vacuole showing K13 (green) with (A) KAE, ***B***. AP-2α, ***C***. AP-2µ, ***D***. AP-1/2β, ***E***. AP-2σ, and ***F***. DrpC (all magenta). The IMC1 (blue) shows the parasite inner membrane complex, and zoomed panels are taken from yellow boxed regions. See Fig S1 for widefield fluorescence images. Scale bars for large and small panels are 5 µm and 0.5 µm, respectively.

Dynamin typically mediates vesicle fission during endocytosis. *T. gondii* has three dynamin-related proteins (Drp) with DrpA and DrpB known to have roles in apicoplast division and vesicle trafficking to rhoptries and micronemes, respectively (Breinich et al., 2009; van Dooren et al., 2009). DrpC (TGME49_270690) function is less well understood although its depletion leads to failure of cell division and proliferation (Melatti et al., 2019). DrpC pulldowns have implicated an association with both AP-2 components and K13 (Heredero-Bermejo et al., 2019) and, when we C-terminally reporter tagged this protein, DrpC is seen associated with the K13 complex (Fig 1F, S1). DrpC is located at a relatively interior position with respect to K13 and appears in one of two forms — a ring of similar diameter to that of K13, or a conical punctum of smaller diameter — consistent with cycles of constriction (Fig 1F, S2C-F).

To identify further molecular components of this conspicuous K13-associated structure in *T. gondii*, we used proximity-dependent biotinylation (BioID) with five bait proteins (K13, KAE, AP-2α, AP-2μ, DrpC). Enrichment of biotinylated proteins (≥2-fold) for each bait compared to a *birA\**-negative control cell line identified proteins as possible K13 complex interactors, with many candidates identified by multiple of the baits (Fig 2A, S3A). Several known proteins of the IMC were identified in the enriched set consistent with the peripheral location of this structure. Twenty identified proteins of previously uncharacterized location were reporter-tagged (Fig 2A, S3) and this validated a further five proteins that exclusively located at the K13 complex (Fig 2B). Four of these proteins are orthologues of *Plasmodium* cytostome proteins (Birnbaum et al., 2020): Eps15-like (Eps15L, TGME49_227800) that contains two Eps15-homology (EH) domains and a Coiled-coil region; UBP1 (TGME49_277895), a putative deubiquitinase that is also implicated in ART-tolerance; metacaspase MCA2 (TGME49_243298); and a <u>C</u>oiled-coil and Gas2-related (<u>GAR</u>) domain protein (CGAR [TGME49_297520], previously named proteophospho-A glycan [PPG] 1 in *T. gondii*). The fifth protein, ISAP1 (TGME49_202220), is specific to coccidian apicomplexans and consists of coiled-coil domains but no other conserved features (Chern et al., 2021). Given the significant overlap of *T. gondii* K13 complex proteins with the *Plasmodium* cytostome we also tested two further orthologues, Kelch13 interaction candidate (KIC) 3 and KIC7, that were identified by BioID of the cytostome although not detected in our BioID results (Birnbaum et al., 2020). KIC7 shares similarity with AGFG (Arf-GAP domain and FG repeat-containing protein) which, in other organisms, has been implicated with Eps15 interaction and endocytosis (Doria et al., 1999; Pryor et al., 2008). The *T. gondii* AGFG orthologue (*Tg*AGFG, TGME49_257070) was exclusively located with the *T. gondii* K13 complex (Fig 2B). The orthologue of KIC3 (TGME49_265420), however, does not localize to the K13 foci in *T. gondii* (Fig. S3B). The reporters for Eps15L, MCA2, CGAR and ISAP1 collocate with K13 as a ring, and UBP1 and AGFG both form a punctum within the K13 ring. These six further K13 complex proteins support that a similar molecular apparatus to the *Plasmodium* cytostome occurs at the periphery of *T. gondii* tachyzoites

**Figure 2.**
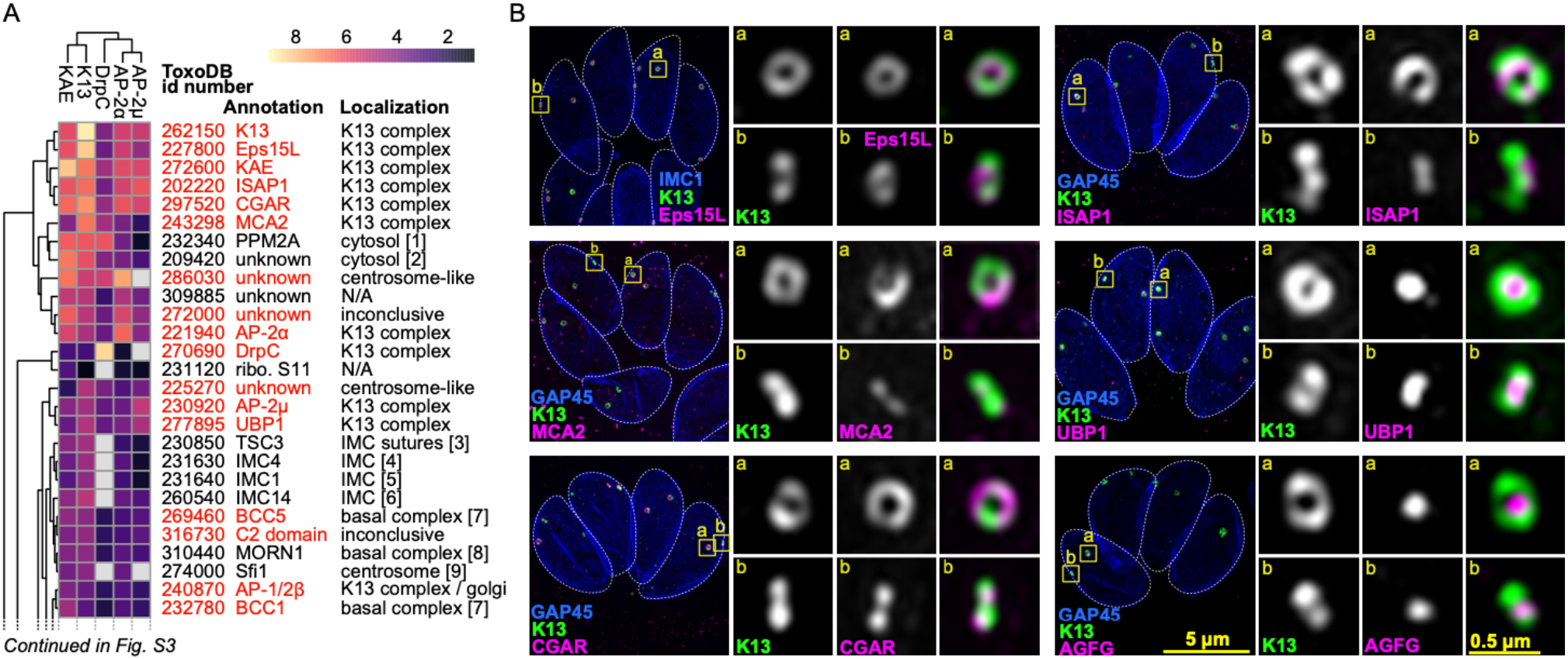
*Toxoplasma* K13 complex shows conservation with the *Plasmodium* cytostome but also unique proteins. ***A***. Heat map of BioID-enriched proteins determined with five bait proteins. Results are clustered according to similarity of results between baits, and ordered by fold enrichment (colors). Proteins reporter-tagged in this study are shown in red, and inferred cell location given. Numbers in brackets refer to previously published localizations: [1] (Yang et al., 2019a), [2] (Seidi et al., 2018), [3] (Chern et al., 2021), [4] (Hu et al., 2006), [5] (Hu et al., 2002), [6] (Anderson-White et al., 2011), [7] (Engelberg et al., 2021), [8] (Gubbels et al., 2006), [9] (Suvorova et al., 2015). Full heat map is shown in Fig S3A. ***B***. Collapsed projections of 3D-SIM images of intracellular parasites showing K13 (green) with Eps15L, MCA2, CGAR, ISAP1, UBP1, and AGFG (all magenta). The IMC1 or GAP45 (blue) mark the parasite pellicle, and zoomed panels are taken from yellow boxed regions. Scale bars for large and small panels are 5 µm and 0.5 µm, respectively.

### The K13 complex is a stable and essential component of the *Toxoplasma* inner membrane complex

The *Plasmodium* blood-stage cytostome occurs in the trophozoite stage where the inner membrane complex (IMC) is absent and, hence, access to the parasite plasma membrane is unencumbered. The plasma membrane of *T. gondii* tachyzoites, on the other hand, is heavily fortified by a robust IMC comprising a dense filamentous proteinaceous network that supports rectangular membranous cisternae sutured together as a quilt adjacent to the cytosolic face of the plasma membrane (Anderson-White et al., 2012; Morrissette & Sibley, 2002). To determine how the K13 complex is positioned with respect to this IMC, and how it might have access to the plasma membrane for possible endocytic function, we co-labelled several components of these peripheral structures with *Tg*K13.

The *Toxoplasma* IMC membrane plate boundaries are demarcated by suture protein ISC3 (Chen et al., 2017). K13 is consistently located at these boundaries (Fig 3A). K13 occurs either within the longitudinal or the transverse plate boundaries, including the most apical boundary at the base of the conically shaped apical cap plate. The rings appeared to be excluded from the intersection of longitudinal/transverse boundaries and, while predominantly in the apical half of the cell, the rings are seemingly haphazardly dispersed in IMC position. Markers for the IMC cisternae outer (GAP45) and inner (GAPM1a) membranes showed that the K13 ring is positioned in windows where these membrane markers are excluded (Fig 3B, C) (Harding et al., 2019). Such mutual exclusion is also seen for K13 and the proteinaceous network marker IMC1 (Fig 3D) (Anderson-White et al., 2012). Together these data indicate coordinated interruption of the IMC where the K13 complex provides an interface between the plasma membrane and the cell cytoplasm. The outer leaflet of the plasma membrane is dominated by the GPI-anchored surface antigen glycoprotein SAG1. When SAG1 is immuno-labelled conspicuous foci are seen in the otherwise uniform SAG1 cell-surface distribution, and these foci locate directly above the center of the K13 rings (Fig 3E). This SAG1 pattern suggests either a depression or some other coordination of the plasma membrane with respect to the K13 complex, evidence of an interaction between these cell structures.

**Figure 3.**
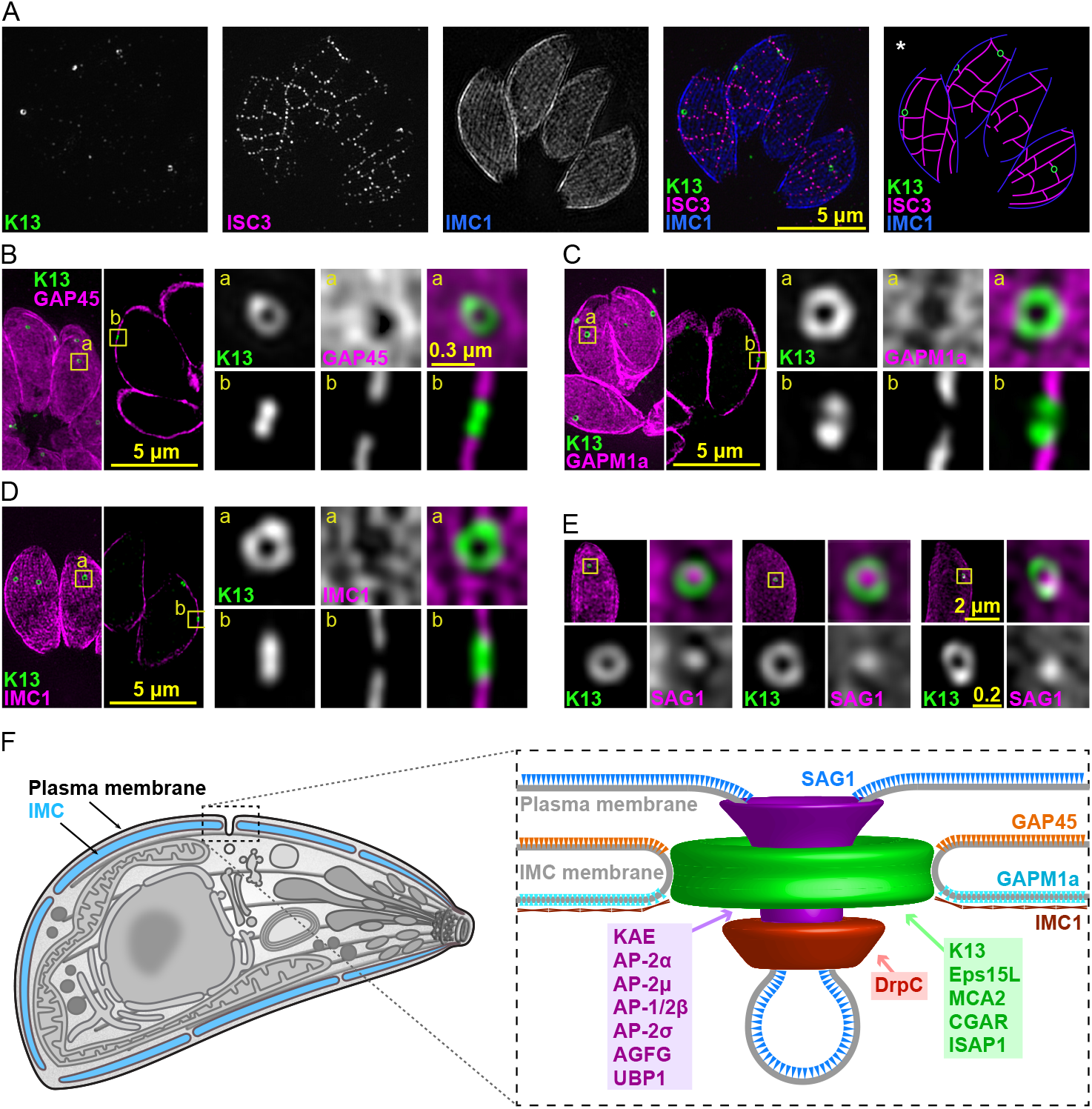
The K13 complex occurs in openings of the IMC at the alveolar plate boundaries. SIM images of intracellular parasites showing K13 (green) with: ***A***. suture protein ISC3; ***B***. IMC outer and ***C***. and inner membrane proteins GAP45 and GAPM1a, respectively; ***D***. IMC subpellicular network protein IMC1; and ***E***. GPI-anchored surface protein SAG1 (all in magenta). Yellow squares correspond to magnified panels, and all scale bars are in µms. ***F***. Model of the endocytic K13 complex in the pellicle of *T. gondii* tachyzoite from these data and Figures 1 and 2. IMC, inner membrane complex.

Endocytosis is typically a dynamic process with many of the molecules that drive it recruited then released after endocytic events and in response to multiple signaling networks (Beacham et al., 2019; Mettlen et al., 2018; Taylor et al., 2011). To test if the K13 endocytic complex shows this typical dynamism, we monitored the locations of K13 and DrpC over time using live-cell fluorescence microscopy. We saw no evidence of temporary recruitment or release of these proteins, and the number and relative positions of the complexes was unchanged over at least 48 min (Fig 4A). These time-lapse data suggest stable structures permanently embedded within the IMC pellicle, and this raised the question of when these structures are first assembled. During the division of most apicomplexan cell stages the daughter cell IMC is prefabricated as internal cups within the mother cell (Anderson-White et al., 2012; Francia & Striepen, 2014). These IMC cups consist of both the membranous cisternae and proteinaceous network elements, and they develop to a state of relative maturity before organelles are sorted into the base of each one. Only during cytokinesis is the mother cell’s plasma membrane recruited to each emerging daughter. To determine how the development of the IMC accommodates the K13 complex, we examined the location of K13 complex components during the early assembly of daughter cells. We saw K13, KAE, AP-2α, EPS15L and DrpC all present as the characteristic complexes in these developing daughter IMCs (Fig 4B). In some images K13 complex components are seen recruited at the growing margin of the IMC1 cup suggesting that this complex is part of the initial assembly of the IMC structure. It is note-worthy that these proteins are all recruited before the plasma membrane is associated with the IMC that ultimately occurs with daughter cell emergence.

**Figure 4.**
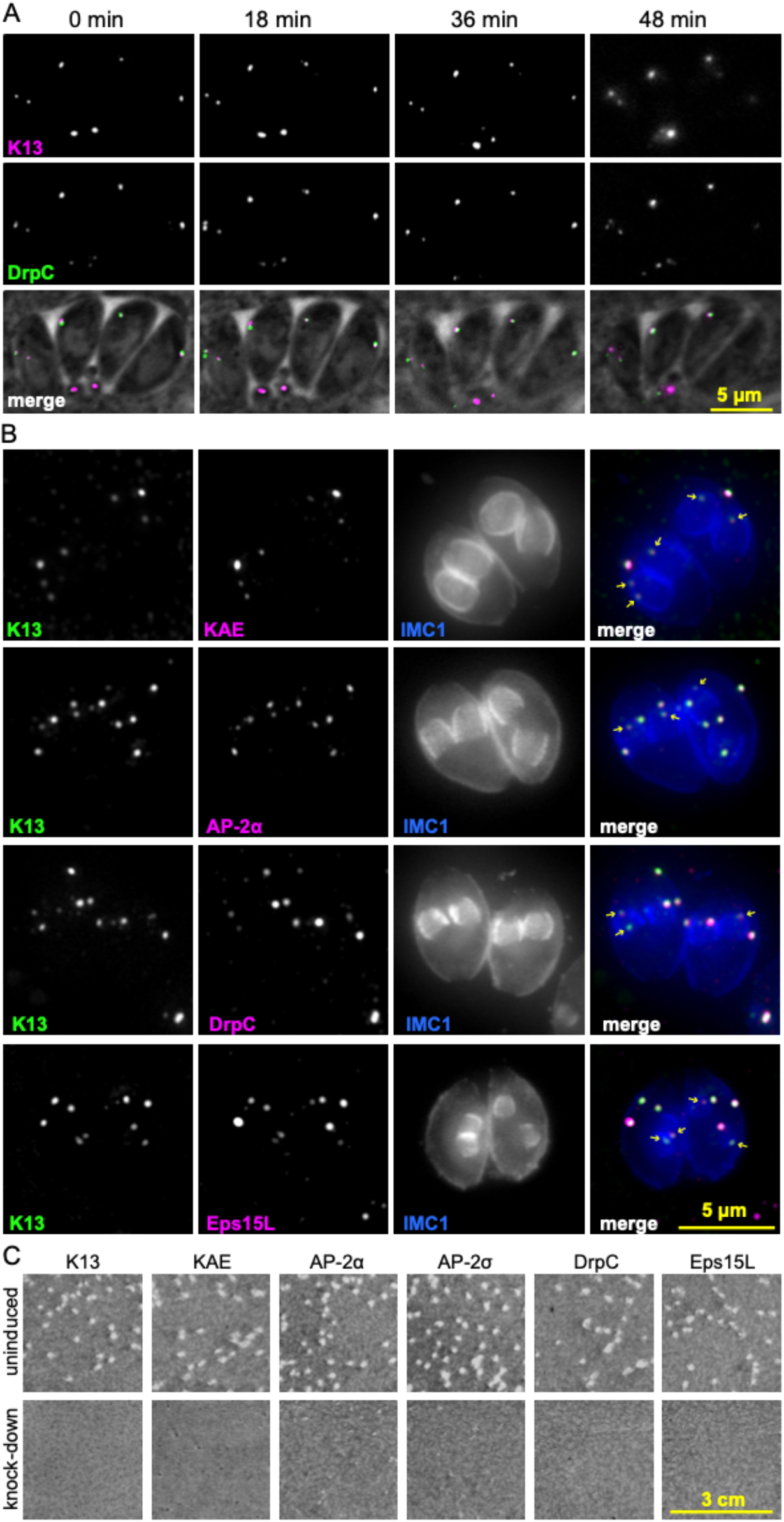
The K13 complex is stable over time, is assembled early in IMC formation during cell replication, and is essential for parasite proliferation. ***A***. Live-cell time-lapse sequence showing four parasites in a host vacuole and that K13 and DrpC locations are stable over time. ***B***. Wide-field fluorescence images of cells fixed during the formation of daughter cells that are evident by internal IMC1 (blue) pellicle ‘cups’. K13 (green) is co-stained with KAE, AP-2α, DrpC and Eps15L (all magenta) with weaker protein signals present for all in the developing daughters (arrows), some of which are seen at the pellicle leading edges. ***C***. Seven-day plaque growth assays in six cell lines with K13 complex gene expression individually suppressed by ATc.

To test if the components of the K13 complex are necessary for tachyzoite growth, seven-day plaque growth assays were performed in host cell monolayers in cell lines engineered for inducible protein knockdowns using an ATc-responsive promoter. We targeted six proteins spanning the different functional components of the complex for individual depletion (K13, KAE, AP-2α, AP-2σ, DrpC and Eps15L) and all mutants showed strong growth phenotypes (Fig 4C).

### Development of a new endocytosis assay for intracellular *Toxoplasma* tachyzoites

Given the composition of the *T. gondii* K13 complex includes several endocytosis-related proteins, we sought to verify that endocytosis is associated with this structure in *T. gondii* tachyzoites. It has been previously shown that fluorescent reporter proteins expressed in the cytoplasm of parasitized host cells can accumulate in the *T. gondii* ‘plant-like vacuole’ or VAC (McGovern et al., 2018). These signals are only seen when proteolytic digestion by the VAC cathepsin L is inhibited, consistent with this compartment functioning as terminal lysosome. Although insight was recently gained for how host proteins traverse the parasitophorous vacuole membrane in which the parasites reside (Rivera-Cuevas et al., 2021), how the material crosses the parasite plasma membrane is uncharacterized. Nevertheless, internalization presumably involves an endocytic process at the parasite surface, and this system offered the opportunity to assay for this activity. We tested if K13 depletion in tachyzoites led to a detectable difference in VAC-accumulation of host reporter protein. K13 was depleted first for 48 hours in wildtype host cells (Fig S4A), and these cells were then allowed to invade host cells expressing cytosolic mCherry. Here they were allowed to grow for 24 hours with continued K13 suppression and cathepsin L inhibition. This assay detected no difference between K13 depletion and controls (Fig S4B). However, in either treatment only approximately 5% of parasites showed detectable accumulation of mCherry indicating that an alternative assay was required to better discern any effects on endocytosis in the knockdowns.

The *T. gondii* surface protein SAG1 is recycled through endocytosis in extracellular tachyzoites (Gras et al., 2019) so we targeted this molecule to develop a new endocytosis assay for parasites within the host cell vacuole. Previous assays for endocytosis of SAG1 in extracellular parasites used antibodies targeting SAG1 (Gras et al., 2019) but these antibodies are known to also inhibit invasion (Mineo et al., 1993) and, therefore, are incompatible with the analysis of intracellular parasites. Instead, we modified the *sag1* gene to express a fused HaloTag that can catalyze covalent linkage to exogenous fluorescent ligands. We used two ligands, one membrane impermeable (HaloTag-Alexa Fluor-660 ligand) and one membrane permeable (HaloTag-Oregon Green) to differentially label the external pool of SAG1 (at the plasma membrane, PM-SAG1) from internal pools of SAG1 (internal SAG1-containing vesicles, Int-SAG1) (Fig 5A). Application of the membrane impermeable ligand first was shown to saturate binding to the PM-SAG1 pool such that secondary labelling with membrane permeable ligands exclusively labelled the Int-SAG1 pools (Fig S4C-F).

**Figure 5.**
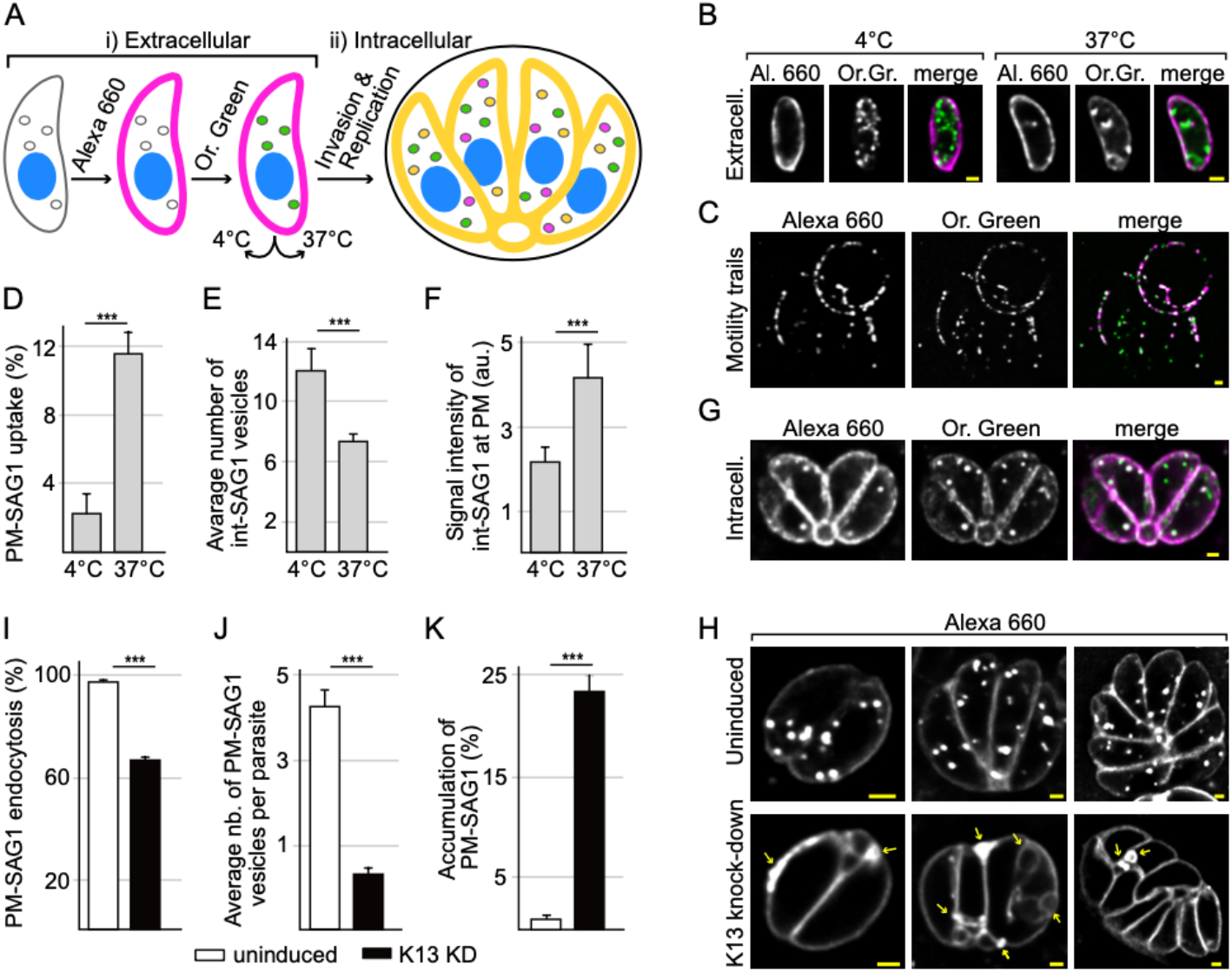
SAG1-Halo bound ligands report on endocytosis which is reduced in K13-depleted parasites. ***A***. Schematic of differential labelling of SAG1 pools at the plasma membrane surface (PM-SAG1 stained with Alexa 660, magenta) or in internal vesicles (Int-SAG1 stained with Oregon Green, green) and tests for endocytic recycling at either non-permissive (4°C) or permissive (37°C) temperatures (i). Infection of hosts after labelling allows these processes to be monitored during intracellular growth (ii) (orange indicates co-location of signals). ***B***. Ligand detection of labelled extracellular cells incubated at either 4°C or 37°C. ***C***. Motility trails of labelled SAG1 left by gliding parasites at 37°C. ***D-F***. Differences according to temperature treatment in: (***D***) the percentage of cells with internalized PM-SAG1 as a measure of endocytosis, (***E***) the number of Int-SAG1 vesicles per cell as a measure of exocytosis, and (***F***) the intensity of Int-SAG1 at the cell surface as a further measure of exocytosis. ***G***. Redistribution of differentially labelled extracellular cells (PM-SAG1 and Int-SAG1) viewed 24 h after invasion and intracellular growth. ***H***. Alexa 660-labelled PM-SAG1 after 24 h of growth without or with K13 depletion. ***I***-***J***. Reduced uptake of PM-SAG1 to intracellular vesicles is quantified as (***I***) percentage of parasites with internalized PM-SAG1 and (***J***) the average number of PM-SAG1-positive vesicles per cell. ***K*** K13-depleted cells showed accumulations of extra PM-SAG1-positive membranes (arrows in H) which occurred in a higher percentage of cells than for the uninduced controls. Three biological replicates were used for all analyses; P-values are where 0.0001<P≤0.001, *** All scale bars = 1 µm.

To test if this labelling strategy can detect the endocytic/exocytic cycling of SAG1, we first tested extracellular tachyzoites where uptake of SAG1 was initially reported (Gras et al., 2019). PM- and Int-SAG1 pools were separately labelled and then parasites were incubated for 60 minutes at temperatures either permissive (37°C) or restrictive (4°C) to vesicular transport (Fig 5A). At 37°C internalization of PM-SAG1 (Alexa Fluor-660-labelled) into cytosolic puncta was observed and, as evidence of this being vesicle-mediated, this was significantly higher than for the 4°C treatment (Fig 5B, D). Concurrently, Int-SAG1 (Oregon Green-labelled) was observed redistributed at the cell surface in a temperature-dependent manner and this ligand was also detected in the parasite motility trails that are known to contain sloughed surface SAG1 (Fig 5B, C, Furthermore, fewer Int-SAG1 vesicles were observed per parasite (Fig 5B, E). These data are consistent with both endocytic and exocytic recycling of SAG1 being detectable using ligand-conjugated SAG1-Halo which is equivalent to that seen using antibodies (Gras et al., 2019).

To evaluate endocytic/exocytic activity of intracellular parasites, SAG1 dual stained parasites (PM-SAG1 Alexa 660; Int-SAG1 Oregon Green) were allowed to invade host cells (Fig 5A). After 24 hours of growth and replication, the PM-SAG1 signal was observed in internal vesicles in the parasites of most vacuoles (98.3±1.6 %) and the Int-SAG1 signal was detected at the parasite plasma membrane in all (Fig 5G). These signals show that PM-SAG1 is endocytosed and recycled in intracellular tachyzoites, and that SAG1 from the initial mother cell continues to be recycled to daughter parasites through subsequent cell divisions. SAG1, therefore, provides a very effective marker of endocytosis in intracellular tachyzoites, as well as reporting on exocytosis and recycling of surface molecules including to daughter cells.

### K13 depletion disrupts endocytosis and plasma membrane homeostasis in *Toxoplasma*

To test if the K13 complex is involved in endocytosis, the SAG1-endocytosis assay was employed in K13-depleted cells (iΔHA -*Tg*K13-SAG1-Halo). These parasites were pre-treated with ATc for 48 hours to deplete detectible K13. Parasites were then egressed and PM-SAG1 was labelled with the Alexa 660 ligand before being allowed to reinvade host cells and replicate for a further 24 hours with continued ATc-suppression of K13. Endocytic activity was then determined both by the number of parasites with endocytosed PM-SAG1 vesicles and the average number of these vesicles per parasite. Compared to the controls, both measures of endocytosis were significantly reduced in the K13-depleted cells: 97.2±1.3% versus 66.8±1.2%, and versus 0.6 vesicles, respectively (Fig 5H, I and J). We noted also that the K13-depleted cells showed accumulation of extra PM-SAG1-positive membranes attached to or between the parasites and these irregularities were significantly different to the controls (Fig 5H and K). These data support that K13 is associated with the sites and activity of endocytic recycling of SAG1 in intracellular *T. gondii* tachyzoites.

*Plasmodium* blood-stage parasites are dependent on cytostome-mediated endocytosis of erythrocyte cytoplasm for its nutrition, however, the important roles for endocytosis in *T. gondii* intracellular tachyzoites is poorly understood. If endocytosis is necessary for nutrition and growth, a retardation in cell replication would be expected when this process is disrupted. To test for an effect on replication, we depleted K13 for 48 hours in host cells, and then inoculated these parasites into new host cells with ongoing K13 suppression. No difference was seen in parasite replication rate over 24 hours of further growth in the absence of K13 (Fig 6A). The organization of the parasites within the parasitophorous vacuole, however, was noted to be atypical. Replicating *T. gondii* tachyzoites are tightly organized typically presenting as a rosette in in vitro culture in fibroblast monolayers. These rosettes form by parasites being tethered at their bases to the so-called ‘residual body’ — a membrane-bound structure that connects and physically coordinates replicated cells in a state of incomplete cytokinesis until egress is triggered (Periz et al., 2017). At 24 hours post reinfection, the K13-depleted vacuoles showed an increased frequency of disorder compared to K13-replete cells (Fig 6B), and with further time individual parasites were no longer clearly distinguishable in the vacuoles by transmitted light microscopy (Fig S5). We, therefore, used a live-cell plasma membrane marker, a GFP fusion of the integral membrane carrier protein HP03, to visualize the parasite bounding membrane during K13-depleted parasite development (Sangaré et al., 2016). HP03 location showed an accumulation of excess membrane including extra signals at the base and between the parasites (Fig 6C and Fig S5). Transmission electron microscopy (TEM) of thin sections of parasite vacuoles showed that, rather than parasites being organized around and connected to a discrete residue body (uninduced control), K13-knockdown parasite vacuoles showed inter-parasite spaces filled by single-membrane bound parasite cytoplasm (Fig 6D). These extensions lacked an IMC but often contained recognizable parasite organelles. These extensions are consistent with the accumulation of Halo-tagged SAG1 and HP03-GFP seen between parasites (Fig 5H, 6C and S5). Collectively, these data suggest that a failure of plasma membrane reuptake when endocytosis is inhibited leads to an excess of membrane and breakdown of the organization of tachyzoites in the parasitophorous vacuole.

**Figure 6.**
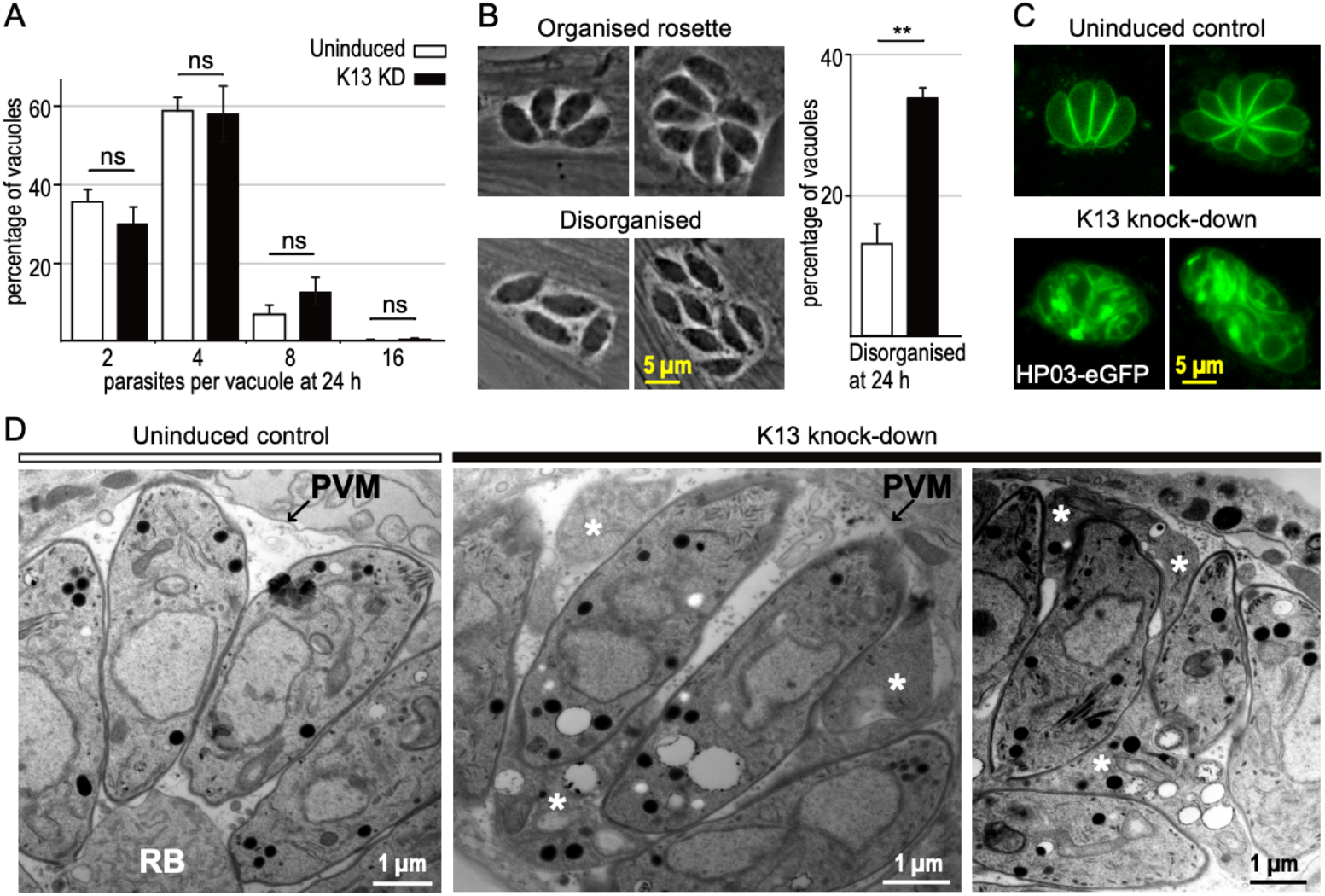
K13-depletion disrupts parasite order and integrity rather than replication rate. ***A***. Number of parasites per parasitophorous vacuole after 24 hours of post invasion replication with or without ATc-induced K13 depletion. ***B***. Percentage of disorganized vacuoles after the same treatments as (A). Disordered vacuoles are scored as those that lack ≻75% of parasites sharing a common posterior orientation as shown by four- and eight-cell parasite rosettes (examples imaged by phase contrast). ***C***. Live cells expressing a plasma membrane marker HP03-eGFP after the same treatments as (A) and (B) showing increasing lack of organization and PM integrity with K13-depletion (see Fig S5 for more examples). Three biological replicates were used for all analyses; P-values are indicated as 0.05<P≤1, ns; 0.01<P≤0.05, *; 0.001<P≤0.01, **; 0.0001<P≤0.001, ***. ***D***. TEM images of parasites within the parasitophorous vacuole with K13 either present or depleted. With K13 depletion the spaces between parasites are filled with parasite cytoplasm containing organelles (asterisks) and bounded by a single membrane. RB, residual body; PVM, parasitophorous vacuole membrane.

### The K13 complex is an ancient endocytic adaptation common to Myzozoa

Given the common presence of a specialized K13-associated endocytic structure in both *Plasmodium* and *Toxoplasma*, we asked how ancient this association is and in what other taxa it might be found. K13, as signature for this structure, is comprised of a highly conserved coiled-coil region and BTB domain attached to the β-propeller kelch domain (Fig 7A). While these domains contribute to a wide range of eukaryotic proteins, the architecture of the K13 protein, including the conserved coiled coil sequence, could only be found within apicomplexans and other members of Myzozoa (chrompodellids, dinoflagellates, perkinsids). Both dinoflagellates and chrompodellids typically have two K13 proteins and the molecular phylogeny of K13 shows that an ancestral gene duplication gave rise to these two paralogues but that apicomplexans lost one of these (Fig 7B). A further conspicuous feature of the K13 complex is the duplication of the AP-2α ear domain to form two proteins — *Tg*AP-2α with a degenerate ear domain, and KAE that contains a conserved C-terminal ear domain but this larger protein otherwise lacks further AP-2α similarity or other conserved domains (Fig 7A, Fig S6). Structural modelling of the ear domain of KAE predicts that it maintains the conserved motif-binding domains for its interaction partners including the sandwich domain that binds WxxF motifs found in proteins such as Eps15 (Fig S7A-C). *Tg*Eps15L is part of the K13 complex (Fig 2), and we verified the predicted interaction by far-western blot interaction assays where the ear domain of KAE bound *Tg*Eps15L in a WxxF motif-dependent manner (Fig 7C). *Tg*AP-2α did not bind *Tg*Eps15L in this assay (whereas the murine AP-2α did) providing strong evidence of a bifurcation of canonical AP-2α function in this endocytic complex. We find the KAE protein throughout myzozoan organisms where K13 is present. Furthermore, an ancient duplication creating two KAE paralogues is maintained only in dinoflagellates and chrompodellids, and this correlates with the duplication of K13 and its subsequent reduction to one paralogue in apicomplexans (Fig 7B). This evolutionary pattern suggests cognate interactions between K13 and KAE. Collectively, these molecular signatures of the K13 complex found throughout Myzozoa indicate common adaptations of endocytosis that evolved early in the evolution of this group and have been maintained throughout its extant diverse members.

**Figure 7.**
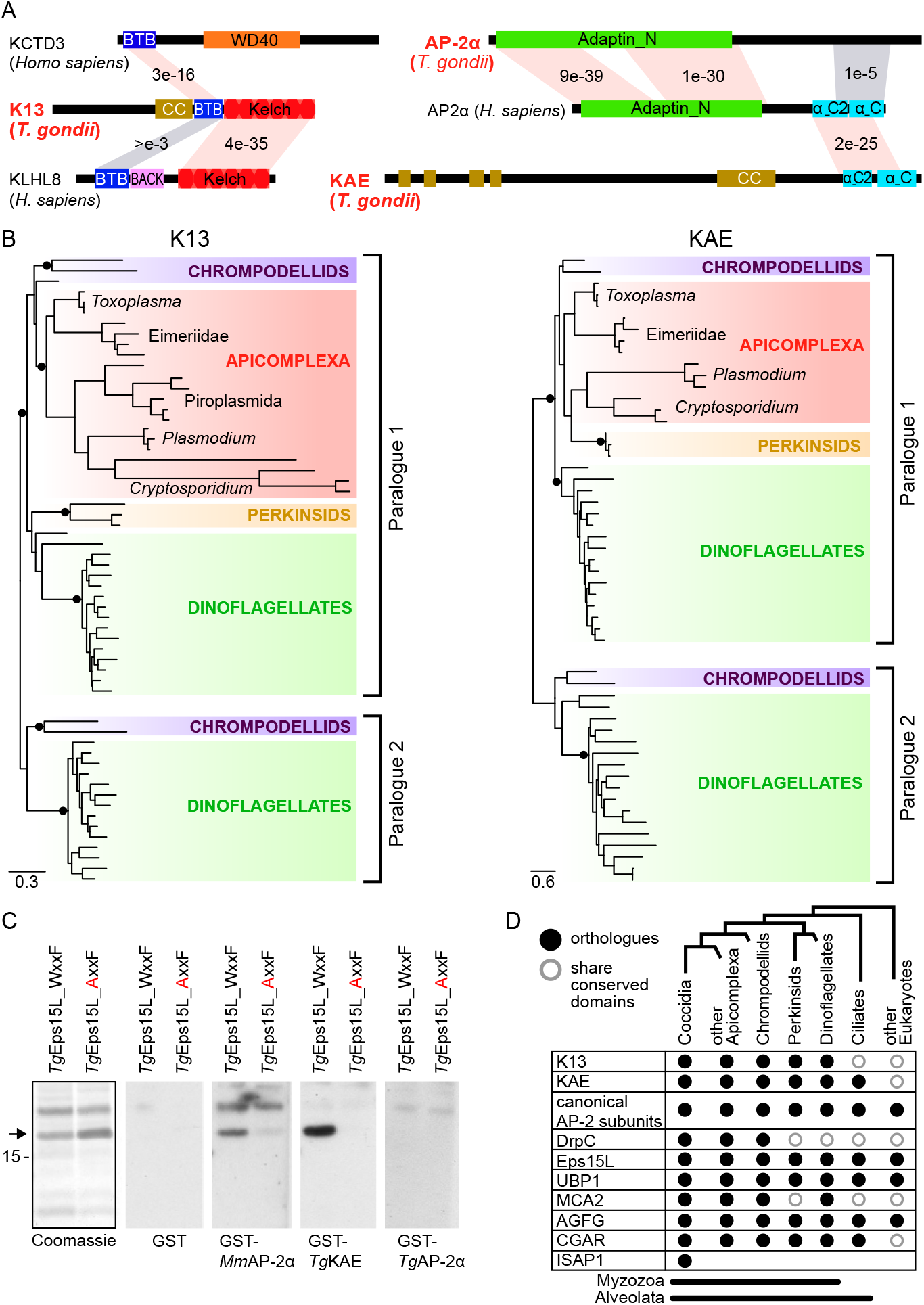
K13 and AP-2 unique traits co-evolved in Myzozoa. ***A***. Domain architectures of: i) K13 compared to proteins from humans, which demonstrate high sequence similarity to the BTB and Kelch domains, and ii) apicomplexan AP-2α and KAE compared to human AP-2α as canonical representative. BLASTP E-values indicate relative conservation between common domains. Conserved domains shown in color; CC, coiled coil. ***B***. Maximum likelihood phylogenies for K13 and KAE. SH-like aLRT branch supports over 0.9 are indicated by black dots, and complete phylogenies are shown in Fig S7D. ***C***. Far-Western blots with immobilized fragments *Tg*Eps15L (arrow) containing either the native (WxxF) or mutated (AxxF) of the predicted AP-2 ear-binding domain (Coomassie-stained gel, black outline). GST-fused ear domains of AP-2 candidate proteins from *Mus musculus* (*Mm*) or *T. gondii* (*Tg*) (or GST alone as negative control) were allowed to bind to the Esp15L fragments and visualized by anti-GST staining. ***D***. Distribution of K13 complex proteins in myzozoan and related lineages.

## DISCUSSION

The recent discovery that endocytosis at the cytostome of *Plasmodium* blood-stage parasites is central to resistance mechanisms against the lead antimalarial artemisinin has focused new attention on this process in *Plasmodium* (Birnbaum et al., 2020; Xie et al., 2020; Yang et al., 2019b). However, it also raised important broader questions in apicomplexan biology. What is the origin of the cytostome structure in *Plasmodium*? Do homologous structures occur in other stages or other apicomplexans? Indeed, what is the site of endocytosis in other pathogen models such as *Toxoplasma*, what are the properties of such sites, and what is the dominant role for endocytosis in this system? While the mechanisms of endocytosis have been extensively studied in select eukaryotic models (e.g., mammals and plants) these processes are poorly understood in other important organisms including apicomplexans.

As evidence of the *Plasmodium* cytostome being derived from a common structure, we have identified the sites of endocytosis in *Toxoplasma gondii* which share at least 11 proteins in common with the cytostome. Five of these are classically associated with endocytosis: Eps15L and the four AP-2 adaptor complex components. Additionally, the Arf-GAP-related AGFG (KIC7 in *Plasmodium*) is implicated in endocytosis in humans where it contains NPF motifs that bind to the EH domains of Eps15 (Doria et al., 1999), and these NPF motifs also occur in the apicomplexan AGFG.

Other components of the endocytic complex found in *Toxoplasma* and *Plasmodium* do not have direct orthologues in characterized endocytic machineries, but do share potential activities with such systems. KAE, with the conserved α ear domain, likely maintains some of the interactions of the AP-2 complex with other endocytic factors. In mammals, ubiquitination plays an important role in regulating endocytosis. Membrane receptor ubiquitination can induce receptor uptake via recruitment of endocytic adaptors, and the adaptors themselves (e.g. Eps15) can be regulated by cycles of ubiquitination and deubiquitination (Haglund and Dikic, 2012). BTB domain-containing proteins recruit Cullin 3 of the Cullin-RING E3 ubiquitin ligases complexes and are implicated in this process, and many of these BTB proteins contain kelch domains for substrate recruitment (e.g. KLHL8, Fig 7A) (Gschweitl et al., 2016). Apicomplexan K13 contains both BTB and kelch domains, and UBP1 is related to the ubiquitin carboxy terminal hydrolase that targets human Eps15 (Savio et al., 2016), suggesting that both K13 and UBP1 might contribute similar ubiquitin-dependent endocytic function in apicomplexans. The presence of dynamin related protein DrpC at the *Toxoplasma* endocytic site is further suggestive of the membrane fission events of endocytosis, and we observed *Tg*DrpC forming either a ring or constricted puncta just below the K13 ring. *Plasmodium* and other apicomplexans share a DrpC orthologue (Fig 7D), although it was not identified in the *Plasmodium* cytostome interactome (Birnbaum et al., 2020). *Tg*DrpC mutants have previously been implicated with cell replication defects including aberrant mitochondrial morphologies (Melatti et al., 2019), and we cannot exclude possible dual *Tg*DrpC functions including in mitochondrial division. We note, however, that reporter-fused *Tg*DrpC location was sensitive to the terminus being labelled (C-terminal fusions were exclusive to K13 locations, N-terminal fusions include additional apparent distribution including mitochondrial proximity). Furthermore, K13 depletion also resulted in aberrant mitochondrial forms extending beyond the cell so these could represent secondary phenotypes (Fig 6D).

Other proteins of the apicomplexan endocytic apparatus distinguish it from canonical systems (Fig 7D). CGAR contains a C-terminal GAR domain that is predicted to bind microtubules and, therefore, might mediate positioning of the endocytic machinery amongst the subpellicular microtubules of apicomplexans. The cysteine protease metacaspase MCA2 is also common to *Plasmodium* and *Toxoplasma* machineries, although its function here is unclear. The apparent non-involvement of clathrin in endocytosis is also common to the apicomplexans. This classic vesicle coat protein that typically interacts with AP-2 and Eps15 in endocytosis was not detected in reciprocal BioID and pulldown experiments in *Plasmodium* (Birnbaum et al., 2020; Henrici et al., 2020), as was also the case for our *Toxoplasma* BioID data. Thus, while the involvement of clathrin with AP-1 mediated post-Golgi trafficking has been retained in apicomplexans, endocytosis is clathrin-independent (Birnbaum et al., 2020; Henrici et al., 2020; Venugopal et al., 2017). In addition to these common derived features of endocytosis, five proteins of the *Plasmodium* cytostome (KIC1, 5, 6, 8 and 9) lacked detectable orthologues in *Toxoplasma*, and ISAP1 is only identifiable in *Toxoplasma* (and other coccidia). Further, while KIC3 does have an orthologue in *Toxoplasma*, we observed this protein in the basal complex (Fig S3B) suggesting a different functional role in *Toxoplasma*. So, there is evidence of further specialization of the endocytic structures between these parasites.

The most unusual feature of the endocytic structures of apicomplexans is their permanence. In most cells endocytic events are transient, and signal-activated recruitment to the plasma membrane of the proteins that drive endocytosis is integral to its regulation and control (Beacham et al., 2019; Mettlen et al., 2018; Taylor et al., 2011). However, we see no evidence of cycles of recruitment and dissociation of proteins to the endocytic structures of *Toxoplasma*. Moreover, the endocytic apparatus is assembled to a very complete state early in daughter cell formation. This occurs well before these nascent daughter structures are in contact with the plasma membrane that only envelops the daughter cells late in cell division (Sheffield and Melton, 1968). The inner membrane complex of apicomplexans is an elaborate structure and its filamentous proteinaceous network that supports the cell, once formed, may not allow the subsequent access and insertion of the endocytic machinery. It is curious, however, that even the dynamin related protein that likely mediates the final steps of membrane fission is present at these very early stages of cell formation. This indicates that DrpC’s recruitment is independent of the membrane upon which it will ultimately act. The *de novo* formation of endocytic structures in daughters means they occur concurrently with those of the mother cell which do not appear to recycle to their daughters. Rather, we see evidence of accumulation of the mother cell structures at the residual body at the base of the interconnected parasites (Fig 4A). It is of note that several of our BioID false positives were for proteins of either the basal complex or centrosomes (Fig S3). This might reflect developmental proximity of these structures either during new cell formation when the new IMC pellicle coalesces at the duplicated centrosomes, or during relocation of mother cell structures to the cell posterior (Anderson-White et al., 2012). K13 in *Plasmodium* is also seen associated with the new cells of the dividing schizont stage, so this preformation of stable endocytic structures during cell division might be a common feature of apicomplexans (Yang et al., 2019b). While we anticipate that endocytosis is likely to be tightly controlled in apicomplexans as for other eukaryotes, the responses of its molecular components to signaling are seemingly quite different to other models.

The supramolecular organization of these endocytic structures in *Toxoplasma* is highly reminiscent of the ‘micropore’ that has been observed in apicomplexans by TEM for over half a century (Nichols et al., 1994; Scholtyseck and Mehlhorn, 1970; Sheffield and Melton, 1968). The micropore is a small invagination of the plasma membrane that is coated by a thin electron dense layer and that penetrates a dense collar that sits within a break in the IMC. The rings formed by K13, Eps15L, MCA2, CGAR and ISAP1 within the sutured boundaries of the IMC alveolae are consistent with this collar, and the funneled arrangement of AP-2 components in its lumen is consistent with these proteins lining the membrane invagination. It has long been speculated that the micropore is the site of endocytosis, however, neither its function nor molecular composition have been determined previously (Nichols et al., 1994). The micropore is observed throughout Apicomplexa as well as in the related alveolates, dinoflagellates and ciliates, and all share the common IMC-type pellicle beneath their plasma membrane (Appleton and Vickerman, 1996; Moore et al., 2008; Mylnikov, 2009; Nilsson and van Deurs, 1983; Perkins, 1976; Schrével et al., 2016). K13 is present in dinoflagellates (and perkinsids and chrompodelids) and the AP-2α/KAE bifurcation is in all alveolates (Fig 7D). The ring-forming K13, Eps15L, MCA2, ISAP1 and CGAR all contain coiled coil domains or cytoskeletal interaction motifs, that likely facilitate protein-protein integrations that might contribute to the stability of this structure, and KAE also contains coiled coil motifs.

Collectively, these data suggest that the micropore is a common endocytic apparatus that evolved in response to the formation of the alveolate cell pellicle. Apicomplexans have seemingly streamlined this apparatus with the loss of the duplicates of K13 and KAE, although the significance of this is unknown. Moreover, ciliates have independently evolved an elaborate phagocytic oral apparatus for prey capture that can be orders of magnitudes larger than the micropore. While any evolutionary link to the micropore for this derived structure is unknown, this subsequent adaptation might explain ciliate’s apparent loss of K13. The *Plasmodium* cytostome, on the other hand, is likely a genuine manifestation of the micropore as strongly suggested by its protein conservation with that of *Toxoplasma*. During the hemoglobin feeding blood-stage, *Plasmodium* disassembles the IMC alleviating it of these obstacles to endocytosis (Ferreira et al., 2021). However, the IMC is reassembled during schizogony and merozoite formation where K13 rings are also seen, and micropores are observed in merozoites by TEM also (Moore and Sinden, 1974; Wichers et al., 2021; Yang et al., 2019b). It is likely that these micropores differentiate into cytostomes upon the invasion of new blood cells and ring-stage formation.

The functional significance of endocytosis likely varies in different apicomplexans and different cell stages. In *Plasmodium*, disruption of K13 and other cytostome proteins, either through natural ART-selected mutations or experimental perturbations, results in decreased hemoglobin uptake and a delay to blood-stage parasite growth (Birnbaum et al., 2020; Yang et al., 2019b). However, no similar immediate growth delay was seen in *Toxoplasma* with K13-depletion. Endocytosis was not completely ablated in these cells according to our SAG1 assay, and this might reflect either some residual K13 protein, or partial endocytic function even in its absence. But in either case, the dominant phenotype of suppressed endocytosis was a chaotic development of these parasites manifested as an expansion of the plasma membrane such that it, and its cytosolic contents, spilled in between cells. Egress of *Toxoplasma* tachyzoites requires completion of cytokinesis by fission of the individual cells’ connections to the residual body, and this process might not be possible in these poorly membrane-delineated cells. Our data point to plasma membrane homeostasis, rather than nutrition, being the more significant function of endocytosis in *Toxoplasma* tachyzoites. Our endocytosis assay shows that surface membrane recycling occurs in the intracellular stages just as it does in the motile extracellular forms (Gras et al., 2019). A deficiency of membrane uptake compared to delivery, through secretion events such as surface molecule delivery and dense granule secretion, would result in the membrane excess that we observe. Membrane recycling could also be the major role for the micropore of *Plasmodium* merozoites. This transmissive stage, from one blood cell to the next, is not otherwise expected to feed but they do remodel their surface molecules during egress and invasion events. Thus, K13 may contribute to multiple endocytic modalities in *Plasmodium*. Similarly, we cannot eliminate a possible nutritional function for endocytosis in *Toxoplasma*. The observed lower activity of endocytosis of host cytoplasm compared to parasite surface molecules might indicate that it fulfils an alternative role to general nutrition, or its nutritional contribution might be below the threshold of detection in our growth assays. Nevertheless, our identification of a broad range of the molecules of endocytosis in apicomplexans, and the development of an effective assay for endocytosis in intracellular stages, provides wide new opportunities to dissect the processes and purposes of this neglected and evidently divergent cell function in these important parasites.

## ACKNOWLEDGEMENTS

This work was supported by the Wellcome Trust, United Kingdom, Investigator award 214298/Z/18/Z to RFW, and a Deutsche Forschungsgemeinschaft (DFG, German Research Foundation) grant 464878930 to SG. BNMS was supported by a Gates Cambridge Scholarship, and KB was supported by the Leverhulme Early Career Fellowship (ECF-2015-562) provided jointly by the Isaac Newton Trust and the Leverhulme Trust. MSR and JH was supported by Wellcome Trust grants 086598 and 214272 to MSR and a Wellcome Trust Strategic Award 100140 to the Cambridge Institute for Medical Research (CIMR). NRZ was supported by Wellcome Trust grant WT 207455/Z/17/Z. CMK was supported by a Canada Vanier Graduate Scholarship and JBD is supported by the Canada Research Chair (Tier II) in Evolutionary Cell Biology and by the Natural Sciences and Engineering Research Council of Canada (RES0043758, RES0046091). Y.R-C. was supported by NIH awards F31AI152297 and T32AI007528. We thank VEuPathDB for their invaluable Informatics Resources, and Christine Hopp, Nicola Hodson, Julian Rayner, Markus Meissner and Mirko Singer for useful discussions, and Cornelia Niemann for assistance with electron microscopy. We dedicate this report to the memory of David Warhurst.

## METHODS

### Growth and generation of transgenic *T. gondii*

*T. gondii* tachyzoites from the strain RH and derived strains, including RH Δku80/TATi (Sheiner et al., 2011), were maintained at 37 °C with 10% CO2 growing in human foreskin fibroblasts (HFFs) cultured in Dulbecco’s Modified Eagle Medium supplemented with 1% heat-inactivated fetal bovine serum, 10 unit ml^−1^ penicillin, and 10 μg ml^−1^ streptomycin, as described elsewhere (Roos et al., 1995). When appropriate for selection, chloramphenicol was used at 20 μM and pyrimethamine at 1 μM. Reporter protein-tagging of endogenous gene loci with reporters 3x HA, 3x v5 and eGFP was done according to our previous work (Barylyuk et al., 2020). For protein function tests by gene knock-downs, endogenous promoters were replaced with an anhydrotetracycline (ATc)-regulatable t7s4 promoter (Sheiner et al., 2011) using the same strategy as for the endogenous gene fusions. Oligonucleotides used for all gene modifications are shown in Table S1

### Immunofluorescence microscopy

*T. gondii*-infected HFF monolayers grown on glass coverslips were fixed with 2% formaldehyde at room temperature for 15 min, permeabilized with 0.1% TritonX-100 for 10 min, and blocked with 2% BSA for 1 h. The coverslips were then incubated with a primary antibody for 1 h, followed by 1 h incubation with a secondary antibody. Coverslips were mounted using ProLong Diamond Antifade Mountant (ThermoFisher Scientific, Massachusetts, USA). Images were acquired using a Nikon Eclipse Ti widefield microscope with a Nikon objective lens (Plan APO, 100×/1.45 oil) and a Hamamatsu C11440, ORCA Flash 4.0 camera. 3D structured illumination microscopy (3D-SIM) was implemented on a DeltaVision OMX V4 Blaze (GE Healthcare, Issaquah, California, USA) with samples prepared as for widefield immunofluorescence assay (IFA) microscopy with the exception that High Precision coverslips (Marienfeld Superior, No1.5H with a thickness of 170 μm ± 5 μm) were used in cell culture, and Vectashield (Vector Laboratories, Burlingame, California, USA) was used as mounting reagent. Samples were excited using 405, 488, and 594 nm lasers and imaged with a 60x oil immersion lens (1.42 NA). The structured illumination images were reconstructed in softWoRx software version 6.1.3 (Applied Precision). All fluorescence images were processed using Image J software (http://rsbweb.nih.gov./ij/).

### BioID

#### Sample preparation

For the proximity biotinylation assay, we generated 5 different cell lines (in parental line *T. gondii* tachyzoites RH Δku80) by in situ genomic N- or C-terminal tagging of one of the 5 bait proteins (K13 and KAE tagged at the N-terminus, AP-2α, AP-2µ and DrpC all tagged at their C-terminus) with the promiscuous bacterial biotin ligase, BirA*. The parental cell line was used as a negative control in biotin treatments. We followed the BioID protocol according to Chen *et al*. (Chen et al., 2015) and our previous work (Koreny et al., 2021). Briefly, the parasites were grown in the elevated biotin concentration (150 μM) for 24 prior to egress, separated from the host-cell debris and washed 5× in phosphate-buffered saline. The cell pellets were lysed in RIPA buffer by sonication and the lysates containing ∼5mg of total protein content were incubated with 250 µl of unsettled Pierce™ Streptavidin Magnetic Beads (Thermo-Fisher: 88817) overnight at 4°C with gentle agitation. The beads were then sequentially treated as follows: washed 3× in RIPA, 1× in 2 M UREA and 100 mM triethylammonium bicarbonate (TEAB; Sigma); washed 3× in RIPA; reduced in 10 mM DTT and 100 mM TAEB for 30 min at 56°C; alkylated in 55 mM iodoacetamide 100 mM TAEB for 45 min at room temperature in the dark; and washed in 10 mM DTT 100 mM TAEB, followed by 2x 15 min in 100 mM TAEB with gentle agitation. The peptides were digested on the beads by a 1 h 37°C incubation in 1 μg of trypsin dissolved in 100 mM TAEB, followed by an overnight 37°C incubation after adding an extra 1 μg of trypsin.

#### TMT labelling

The peptide concentrations were measured using the Pierce™ Quantitative Fluorometric Peptide Assay (ThermoFisher: 23290) according to the manufacturer’s instructions and 5 μg of the digested peptides were subjected to the tandem mass tag (TMT) labelling using TMT10plex isobaric tagging reagent set (ThermoFisher: 90110). Each TMT reagent vial containing 0.8 mg of the labelling reagents was brought to room temperature and dissolved in 60 µl of LCMS-grade acetonitrile immediately before use. The TMT reagents were then split to three sets without exceeding the labelling capacity, and 20 µl of the TMT reagents were added to each peptide sample. After incubating for 1 hour at room temperature, 8 μl of 5% hydroxylamine (v/v) was added to each sample, followed by incubation for 15 mins to quench the reaction. The TMT-labelled fractions were combined and dried in a vacuum centrifuge (Labconco) at 4°C.

#### Analysis of TMT-labelled peptides by liquid chromatography and tandem mass spectrometry

LCMS analyses were carried out on an Orbitrap™ Fusion™ Lumos™ Tribrid™ mass spectrometer coupled on-line with a Dionex Ultimate™ 3000 RSLCnano system (Thermo Fisher Scientific) as previously described (Barylyuk et al., 2020). The XCalibur v3.0.63 software was used to control the instrument parameters and operation, and record and manage the raw data. The LCMS system was operated in the positive-ion data-dependent acquisition mode with the SPS-MS^3^ acquisition method with a total run time of 120 min. The dried TMT10plex-labelled peptide samples resolubilized in an LC-MS sample loading solution (0.1% aqueous formic acid) at a concentration of approximately 1 µg/µl. Approximately 1 µg of the sample was loaded onto a micro-precolumn (C18 PepMap 100, 300 µm i.d. × 5 mm, 5 µm particle size, 100 Å pore size, Thermo Fisher Scientific) with the sample loading solution for 3 min. Following the loading step, the valve was switched to the inject position, and the peptides were fractionated on an analytical Proxeon EASY-Spray column (PepMap, RSLC C18, 50 cm × 75 µm i.d., 2 µm particle size, 100 Å pore size, Thermo Fisher Scientific) using a linear gradient of 2-40 % (vol.) acetonitrile in aqueous 0.1% formic acid applied at a flow rate of 300 nl/min for 95 min, followed by a wash step (70% acetonitrile in 0.1% aqueous formic acid for 5 min) and a re-equilibration step. Peptide ions were analyzed in the Orbitrap at a resolution of 120,000 in an m/z range of 380-1,500 with a maximum ion injection time of 50 ms and an AGC target of 4E5 (MS^1^ scan). Precursor ions with the charge states of 2-7 and the intensity above 5,000 were isolated in the quadrupole set to 0.7 m/z transmission window and fragmented in the linear ion trap via collision-induced dissociation (CID) at a 35% normalized collision energy, a maximum ion accumulation time of 50 ms and an AGC target of 1E4 (MS^2^ scan). The selected and fragmented precursors were dynamically excluded for 70 s. Synchronous precursor selection (SPS) was applied to co-isolate ten MS^2^-fragments in the linear ion trap with an isolation window of 0.7 m/z in the range of m/z 400-1,200, excluding the parent ion and the TMT reporter ion series. The SPS precursors were activated at a normalized collision energy of 65% to induce fragmentation via high-collision energy dissociation (HCD). The product ions were measured in the Orbitrap at a resolution of 50,000 in a detection range of m/z 100-500 with a maximum ion injection time of 86 ms and an AGC of 5E4 (MS^3^ scan). The mass spectrometry proteomics data have been deposited to the ProteomeXchange Consortium via the PRIDE (Perez-Riverol et al., 2022) partner repository with the dataset identifier PXD034193

#### Raw LCMS data processing

The processing of raw LSMS data for peptide and protein identification and quantification was performed with Proteome Discoverer v2.3 (Thermo Fisher Scientific). Raw mass spectra were filtered converted to peak lists by Proteome Discoverer and submitted to a database search using Mascot v2.6.2 search engine (Matrix Science) against the protein sequences of *Homo sapience* (93,609 entries retrieved from UniProt on 09.04.2018), *Bos taurus* (24,148 entries retrieved from UniProt on 17.04.2017), and *Toxoplasma gondii* strain ME49 (8,322 entries retrieved from ToxoDB.org release 36 on 19.02.2018) (Amos et al., 2022). Common contaminant proteins – e.g., human keratins, bovine serum albumin, porcine trypsin – from the common Repository of Adventitious Proteins (cRAP, 123 entries, adapted from https://www.thegpm.org/crap/) were added to the database, as well as the sequence of the BirA* used to generate the BioID bait proteins by gene fusion. The precursor and fragment mass tolerances were set to 10 ppm and 0.8 Da, respectively. The enzyme was set to trypsin with up to two missed cleavages allowed. Carbamidomethylation of cysteine was set as a static modification. The dynamic modifications were TMT6plex at the peptide N-terminus and side chains of lysine, serine, and threonine, oxidation of methionine, deamidation of asparagine and glutamine, and biotinylation of the peptide N-terminus or lysine side chain. The false discovery rate of peptide-to-spectrum matches (PSMs) was validated by Percolator v3.02.1 (The et al., 2016) and only high-confidence peptides (FDR threshold 1%) of a minimum length of 6 amino acid residues were used for protein identification. Strict parsimony was applied for protein grouping.

TMT reporter ion abundances were obtained in Proteome Discoverer using the most confident centroid method for peak integration with 20 p.p.m. tolerance window. The isotopic impurity correction as per the manufacturer’s specification was applied. For protein quantification, PSMs with precursor co-isolation interference above 50% were discarded, and the TMT reporter ion abundances determined for unique (sequence found in proteins belonging to a single protein group) and razor (if sequence is shared by protein belonging to multiple protein groups, the quantification result is attributed to the best-associated Master Protein) peptides were summed.

#### Statistical analysis of protein enrichment in BioID

Data analysis was performed with *R* v3.6.1 (R Core Team, 2021) using packages *tidyverse* v1.2.1 (Wickham et al., 2019)for data import, management, and manipulation, *Bioconductor* (Gentleman et al., 2004)packages *MSnbase* v2.10.1 (Gatto and Lilley, 2012)for managing quantitative proteomics data, *biobroom* v1.16.0 (https://github.com/StoreyLab/biobroom) for converting *Bioconductor* objects into *tidy data frames*, and *limma* v3.40.6 (Ritchie et al., 2015)for linear modelling and statistical data evaluation.

The protein-level report generated by Proteome Discoverer was imported into R and filtered to remove non-*Toxoplasma* and low-confidence (protein FDR confidence level “Low”, *q* ≥ 0.05). Only Master Proteins with a complete set of TMT abundance values across three replicates of the BioID bait and control samples were considered for the analysis. The protein abundance values in each biological sample were corrected for the total amount using normalisation factors derived from the abundances of two proteins, acetyl-CoA carboxylase ACC1 (TGME49_221320) and pyruvate carboxylase (TGME49_284190). Both proteins are highly expressed, endogenously biotinylated, and reside in the matrix of subcellular compartments, the apicoplast and mitochondrion for ACC1 and PC, respectively, where they are not accessible to the BirA*-fused BioID baits. Hence, these two proteins could serve as suitable internal standards. The normalised protein abundances were log2-transformed and modelled as a simple linear relationship between the abundances described by a constant factor (intercept), which is of no interest to us, and the condition (BirA*-tagged vs. control) using *limma*. If the condition parameter estimated by *limma* linear model was significantly different from zero we concluded that the condition (presence of the BirA*-fused bait) had a significant effect on the protein abundance. Also, *limma* estimated the model parameters taking into account the relationship between protein average intensities and the variance (low-abundance proteins tend to have a greater variance) by empirical Bayesian shrinking of the standard errors towards the trend. This enabled a better control of false discoveries and outliers affording more robust identification of significantly enriched proteins. The resulting *p*-values were adjusted for multiple testing using the Benjamini-Hochberg method (FDR < 1 %). Proteins with the adjusted *p*-value below 0.01 were deemed significantly changing abundance in the BioID bait condition vs. the control, and those of them whose abundance in the BioID bait condition was more than two-fold greater than in the control were considered significantly enriched.

### Plaque assay

To test lytic cycle competence of knock-down cell lines by plaque formation in HFF monolayers, 500 freshly lysed parasites were added to T25 flasks containing HFF monolayer. 0.5 μg/ml of ATc was added to induce the gene knock-down, or omitted for uninduced controls. After 7 days of growth, flasks were aspirated, washed once with PBS, fixed with 5 ml of 100% methanol for 5 mins and stained with 5 ml of crystal violet solution for 15 mins. After staining, the crystal violet solution was removed, and the flasks were washed three times with PBS, dried and imaged.

### Replication assay

For replication assays, the parasites were pre-treated with ATc for 48 hours before the egress from the host cell and subsequent invasion of the HFF monolayer growing on coverslips in 6-well plates. Parasites were allowed to grow for further 24 hrs with ATc. Parasitophorous vacuoles were scored containing either 1, 2, 4, 8 or 16 parasites. A minimum of 200 parasitophorous vacuoles were scored for each of the three biological replicates. P-values were calculated with multiple t-tests and corrected for multiple comparisons using the Holm-Sidak method in GraphPad Prism, v8 (GraphPad, California USA).

### Host cytosolic protein uptake assay

Inducible mCherry HeLa cells previously described (Rivera-Cuevas et al., 2021)were seeded into 6-well plates at a density of 1.5 ×10^5^ cells per well. Cytosolic mCherry expression was induced for 4 days by adding 2 µg mL^-1^ of doxycycline each day. Prior to infection of host cells expressing cytosolic reporter, parasites were treated with 0.5 µg mL^-1^ anhydrotetracycline (ATc) or vehicle control for 48 h. Additionally, parasites were treated with 5 µM of the protease inhibitor LHVS for 24 h to inhibit the degradation of material delivered to the VAC. Cells were infected with 1.0 ×10^6^ parasites and allowed to replicate for 24 h in the presence of ATc or vehicle control and LHVS.

Parasites were harvested for analysis by scraping and syringing the infected monolayer on ice followed by filtration and centrifugation at 1,500 g for 10 min at 4°C. Isolated parasites were then treated with a 1 mg mL^-1^ pronase and 0.01% saponin-1X PBS solution for 1 h at 12°C, centrifuged and washed 3x before adding to Cell-Tak coated slides. Extracellular parasites were fixed with 4% paraformaldehyde for 10 min and permeabilized with 0.1% Triton X-100 for 10 min prior to imaging. At least 200 parasites were analyzed per sample and the percentage of parasites positive for mCherry was quantified by dividing the number of parasites containing mCherry signal derived from the host cytosol divided by the total number of parasites analyzed.

### Uptake of parasite surface protein assay

#### Internal tagging of SAG1 (SRS29B TGME49_233460) by transient CRISPR-Cas9 expression

SAG1-Halo and iΔHA -*Tg*K13-SAG1-Halo were generated by tagging SAG1 in the Δku80 and Δku80 iΔHA -*Tg*K13 line using transient Cas9 transfection targeting the following gRNA: TGCAGCCCCGGCAAACTCCAC(GGG). The strain was generated as previously described for Cas9 tagging (Li et al., 2021). Briefly, gRNA oligos were annealed and ligated into the Cas9 vector and verified by sequencing (Eurofins Genomics). Reparation template DNA were generated by amplifying the halo with homology arm (50bp) with SAG1 by PCR using Q5 High-Fidelity DNA Polymerase (New England BioLabs. The repair template was purified using a PCR purification kit (Blirt). Parasite transfection, sorting and screening for positive mutants was done as previously (Li et al., 2021). Briefly, newly released RHDiCreΔku80 or Tet-K13-Ha Δku80 tachyzoites were transfected with the repair template and 10-12 μg of SAG1-Cas9-YFP. The parasites were mechanically egressed 24 to 48 h after transfection, passed through a 3 μm filter, and those transiently expressing Cas9-YFP enriched via FACS 23 (FACSARIA III, BD Biosciences) into 96-well plates (a minimum of 3 events per well). Resultant clonal lines were screened by IFA for SAG1 labelling and integration was confirmed by PCR.

#### Specific labelling of the plasma membrane SAG1

SAG1-Halo strain was labelled with either the non-permeable Alexa 660 (1/1000) or permeable Oregon green (1/4000) dyes (Promega) in cold media for 1h. Parasites were then washed prior to transfer to Ibidi live cell dishes (29 mm) coated with 0.1% poly-L-Lysine as previously described (Gras et al., 2019). After letting the parasite settle for 15 minutes, parasites where imaged live on a Leica DMi8 widefield microscope attached to a Leica DFC9000 GTC camera, using a 100x objective to compare the difference between permeable and non-permeable dye labelling.

#### SAG1 endocytosis assays

For extracellular tachyzoites, the SAG1-Halo strain was labelled with the non-permeable Alexa 660 dye (1/1000) in cold media for 1h. Parasites were wash 3x to remove excess ligand. Parasites were incubated at 4°C or 37°C on the FBS coated live cell dishes (Ibidi live cell dishes, 29 mm) for 1 h prior to live imaging, as described below, for evaluation of internalization of the SAG1 signal. For SAG1 recycling assays of intracellular tachyzoites, parasites were labelled with Alexa 660 as above and then inoculated onto HFF monolayers in Ibidi dishes. Parasites were allowed to replicate for 24 h prior to imaging. For endocytosis assays with K13-depletion, iΔHA -*Tg*K13-SAG1-Halo parasites were first induced for 48h with or without ATC, before mechanical egress, filtering and labelling. Parasites were then allowed to reinfect new HFFs cells, grow for 24 h under the same ATc treatment and were then imaged live, as described below, for evaluation of internalization of the SAG1 signal. Endocytic activity was assessed by the presence of the PM-SAG1 vesicles (non-permeable halo-Alexa Fluor®-660) and the percentage of parasites showing the presence of vesicles was determined. Mean values of three independent experiments ± SD were determined. The mean number of PM-SAG1 vesicles/parasite was determined from the total number of vesicles inside vacuoles divided by the number of parasites in the vacuole. For membrane accumulation, we scored aggregation of PM-SAG1 as signal in the residual body or outside the typical parasite plasma membrane. For these analyses at least 25 vacuoles per replicates were used and mean values of three independent experiments ± SD were determined. All SAG1-Halo images were acquired on a Leica-DMI8, objective 100x with the LasX software (v3.7.4). Imaged were deconvolved using Huygens essential software (v18.04) and batch express processing. Fiji (v1.53c) was used to analyse the picture and all count were made manually.

### Transmission electron microscopy

For iΔHA -*Tg*K13-SAG1-Halo parasites with or without K13 depletion, the parasites were induced for 48 h with or without ATc, released mechanically and filtered prior transfer to Ibidi μ-dishes previously seede with HFF cells. After 24 h of replication the parasites were fixed with 2.5% glutaraldehyde in 0.1 M phosphate buffer pH 7.4. The parasites were washed three times at room temperature with PBS (137 mM NaCl_2_, 2.7 mM KCl, 10 mM Na_2_HPO_4_,M1.8 mM KH_2_PO_4_, pH 7.4) and post-fixed with 1% (w/v) osmium tetroxide for 1 h. Subsequent to washing with PBSMand water, the samples were stained en bloc with 1% (w/v) uranyl acetate in 20% (v/v) acetone for 30 minutes. Samples were dehydrated in a series of graded acetone and embedded in Epon 812 resin. Ultrathin sections (thickness, 60 nm) were cut using a diamond knife on a Reichert Ultracut-E ultramicrotome. Sections were mounted on collodium-coated copper grids, post-stained with lead citrate (80 mM, pH 13) and examined with an EM 912 transmission electron microscope (Zeiss, Oberkochen, Germany) equipped with an integrated OMEGA energy filter operated in the zero-loss mode at 80 kV. Images were acquired using a 2k x 2k slow-scan CCD camera (Tröndle Restlichtverstärkersysteme, Moorenweis, Germany).

### Binding of α ear appendage domains to WxxF motif of *Tg*Eps15L

The construction of a GST fusion of *Mm*AP-2α ear appendage domain has been previously described (Owen et al., 1999) and encompasses amino acids 695–938 of mouse AP2A2. Equivalent domain boundaries for *Tg*KAE (TGME49_272600) and *Tg*AP-2α (TGME49_221940) ear appendage domains were selected based on homology to MmAP-2α. *Tg*KAE (amino acids 1,406-1,672 end) and TgAP-2α (amino acids 1,015-1,348 end) appendage domains were made synthetically and codon optimized for expression in *E. coli*; adding BamH1 and Xho1 sites for *Tg*AP-2α and BamH1 and Sal1 for *Tg*KAE for in-frame cloning into pGEX4T-1. For fusion protein production, the DNA was transformed into BL21(DE3) competent cells for high protein expression and induced mid-log phase with 0.4 mM IPTG 16-20 h at 22°C. Fusion proteins were recovered using Glutathione-Sepharose 4B (GE Healthcare) and eluted from the beads using 30 mM reduced glutathione. His-tagged fragment of *Tg*Eps15L (TGME49_227800) encompassing amino acids 897-1023 (952-WxxF), whose boundaries were predicted using DomPRED (URL:http://bioinf.cs.ucl.ac.uk/software.html) was made synthetically and codon optimized for expression in *E. coli*. The fragment was cloned using BamH1 and EcoR1 (adding a stop codon) into pTrcHisA. For fusion protein production, the DNA was transformed into BL21(DE3) competent cells (Invitrogen), induced mid-log phase with 0.4 mM IPTG, purified using Ni-nitrilotriacetic acid agarose beads (QIAGEN) and eluted with 250 mM imidazole. Point mutation W952A (952-AxxF) was made using site-directed mutagenesis.

For the far-Western blot assay, 2.5 μg of the His-tagged fusion proteins were subjected to SDS-PAGE on 16% Bolt Bis-Tris gel (ThermoFisher) in MOPS running buffer gels and blotted onto a nitrocellulose membrane. The blot was blocked with 20 mM Tris-HCl, pH 7.5, 150 mM NaCl, 0.05% Tween 20, 0.5% BSA, 3 μM reduced glutathione for 30 min and this buffer was used in all the following steps. The blot was incubated with 10 nM GST, GST-MmAP-2α, GST-TgAPale or GST-TgAP-2α for 1h, washed for 30 min, and then labeled with 1:30,000 anti-GST followed by 1:10,000 anti-rabbit-HRP and developed using ECL Prime (GE Healthcare) and X-ray film.

### Structural Predictions

The 3D structure of *Tg*KAE ear appendage domain was modelled using SWISS-MODEL (Waterhouse et al., 2018) by using as template the mouse Alpha-adaptin Appendage Domain (PDB 1w80; chain A, sequence identity 31%). Using the positions of the bound peptides (FxDxF and WxxF) in 1w80, a resultant *Tg*KAE peptide complex structure was rebuilt and further energy minimized in coot (Emsley and Cowtan, 2004). Figures for the protein structures were drawn with Pymol (DeLano, 2020), and the protein-protein interaction networks were generated with Ligplot+ (Laskowski and Swindells, 2011).

### Homology searching, domain detection and phylogenetic analyses

Protein homologues were searched with the iterative version of the profile hidden Markov models (HMMs) search engine (jackhammer) (Johnson et al., 2010). Conserved domains were detected with InterProScan (Quevillon et al., 2005)and the similarity levels between the *T. gondii* and human BTB, Kelch, Adaptin_N and α_C2/αC (ear) domains were evaluated using a pairwise protein BLAST (https://blast.ncbi.nlm.nih.gov/Blast.cgi). For the phylogenetic analyses, sequences were aligned using Mafft v7.407 with the L-INS-i algorithm (Katoh and Standley, 2013). Phylogenetic analyses were performed locally and using the CIPRES online portal (Miller et al., 2010). Alignments were masked and trimmed manually using Mesquite v3.6 (https://www.mesquiteproject.org) or Jalview (https://www.jalview.org). Bayesian analysis was carried out using MrBayes v3.2.7a35 hosted on CIPRES. Datasets were run under a mixed model with four independent runs of four chains each, sampling every 500 generations up to a total of either 1,000,000 or 10,000,000 MCMC generations, depending on convergence criteria. Trees were summarized discarding the first 20% of samples as burn-in; convergence was assessed by manually expecting the generation vs. log-likelihood plots for stationarity, as well checking parameter PSRF values for various parameters. Maximum-likelihood (ML) rapid boostrapping was carried out using RAxML v8.2.1232 (LG+**r**model, rapid bootstrapping) hosted on CIPRES or locally using PhyML-3.1 or IQ-TREE v1.6.12 (1000 bootstraps, with the optimal substitution model) (Guindon et al., 2010; Nguyen et al., 2015). SH-like aLRT and aBayes branch support values were obtained in PhyML-3.1.

## SUPPLEMENTARY MATERIALS

**Supplementary Table 1**

Oligonucleotides used for genetic modifications

**Figure S1:**
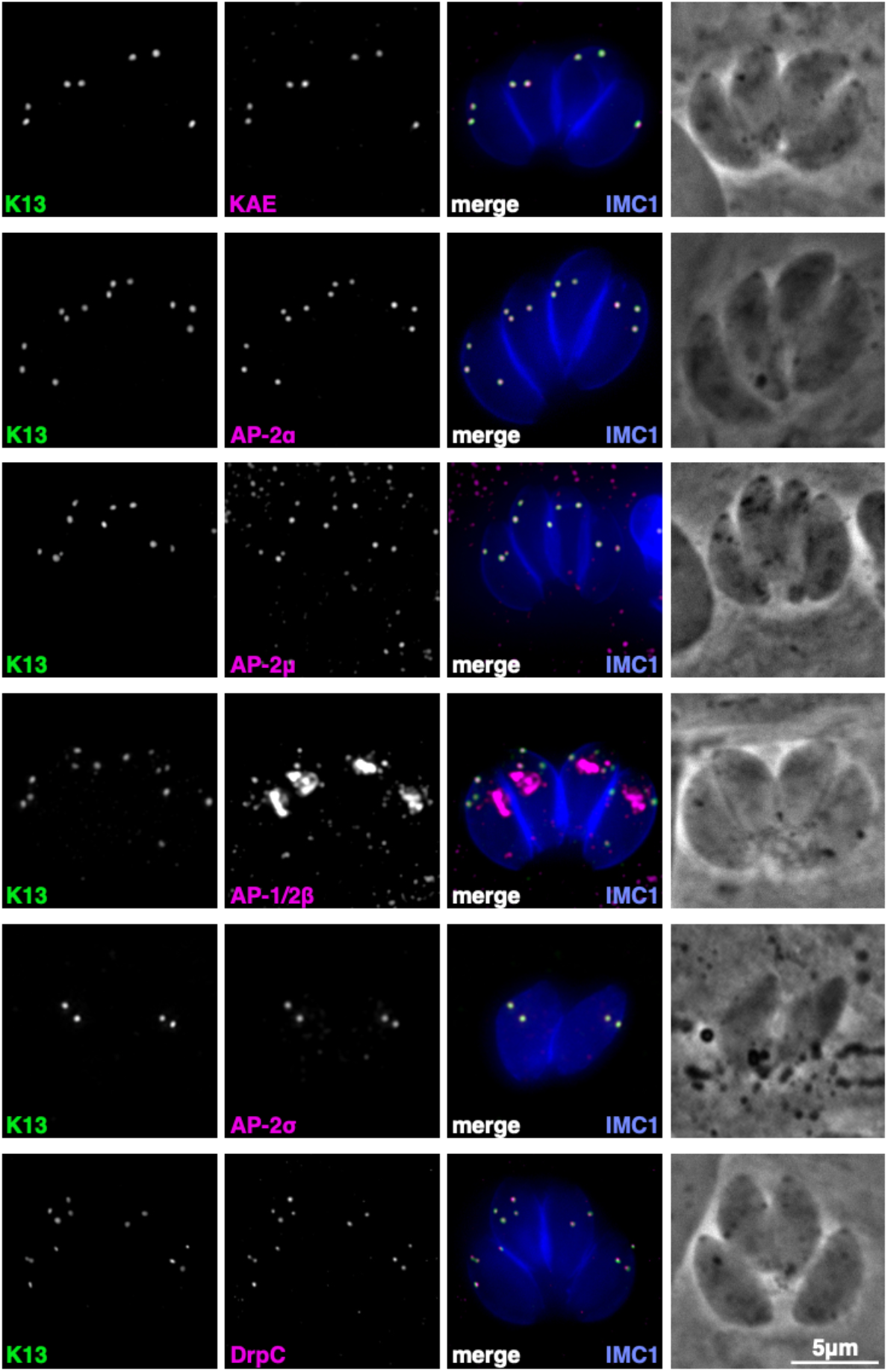
Wide-field fluorescence images of AP-2 subunits, novel AP-2 derived KAE, and DrpC associated with the K13 complex. Only AP1/2β shows additional locations to K13 and this is consistent with it being shared between the AP-2 and the post-Golgi trafficking factor AP-1.

**Figure S2:**
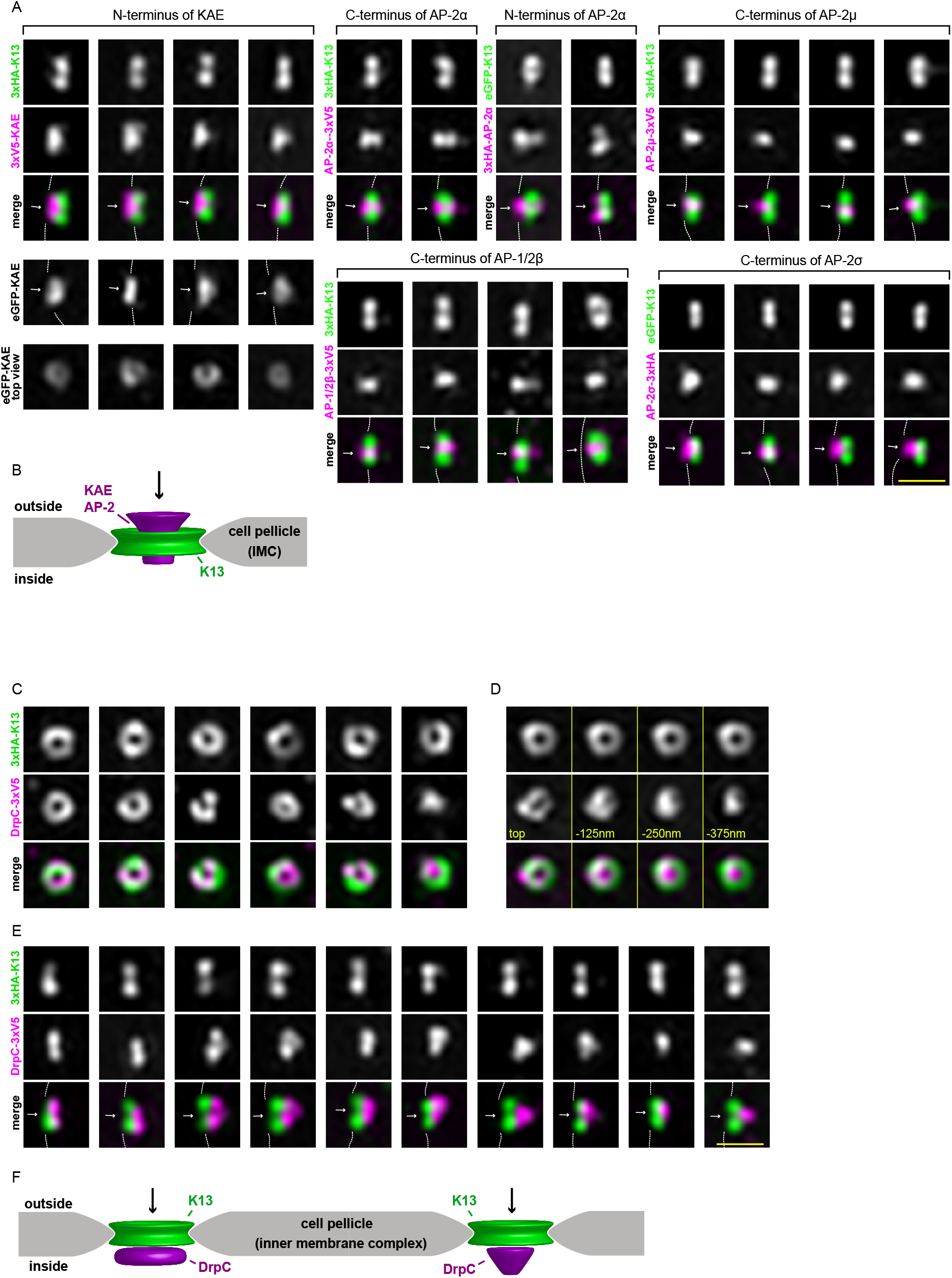
Projections of AP-2, AP-2-derived protein KAE, and DrpC observed by 3D-SIM. (A) Side projections of 3D-SIM images of K13 and the five AP-2 subunits and the AP-2α-derived protein KAE. Arrows indicate the plasma membrane side of the complex, and cross sections of the top of the complex for AP-2ale shows the annular mouth of this funnel. The scale bar for all is 0.5 µm. (B) Model of the AP-2 complex locations with respect to K13 and the IMC. (C) Multiple top projection examples of DrpC states ranging from ring to punctum while K13 rings stay uniform in diameter. (D) Sequence of optical sections of the DrpC signal through an individual K13-DrpC complex. (E) Multiple side projection examples of DrpC states compared to K13. Scale bar for all is 0.5 µm. (F) Model of the AP-2 complex locations with respect to K13 and the IMC.

**Figure S3:**
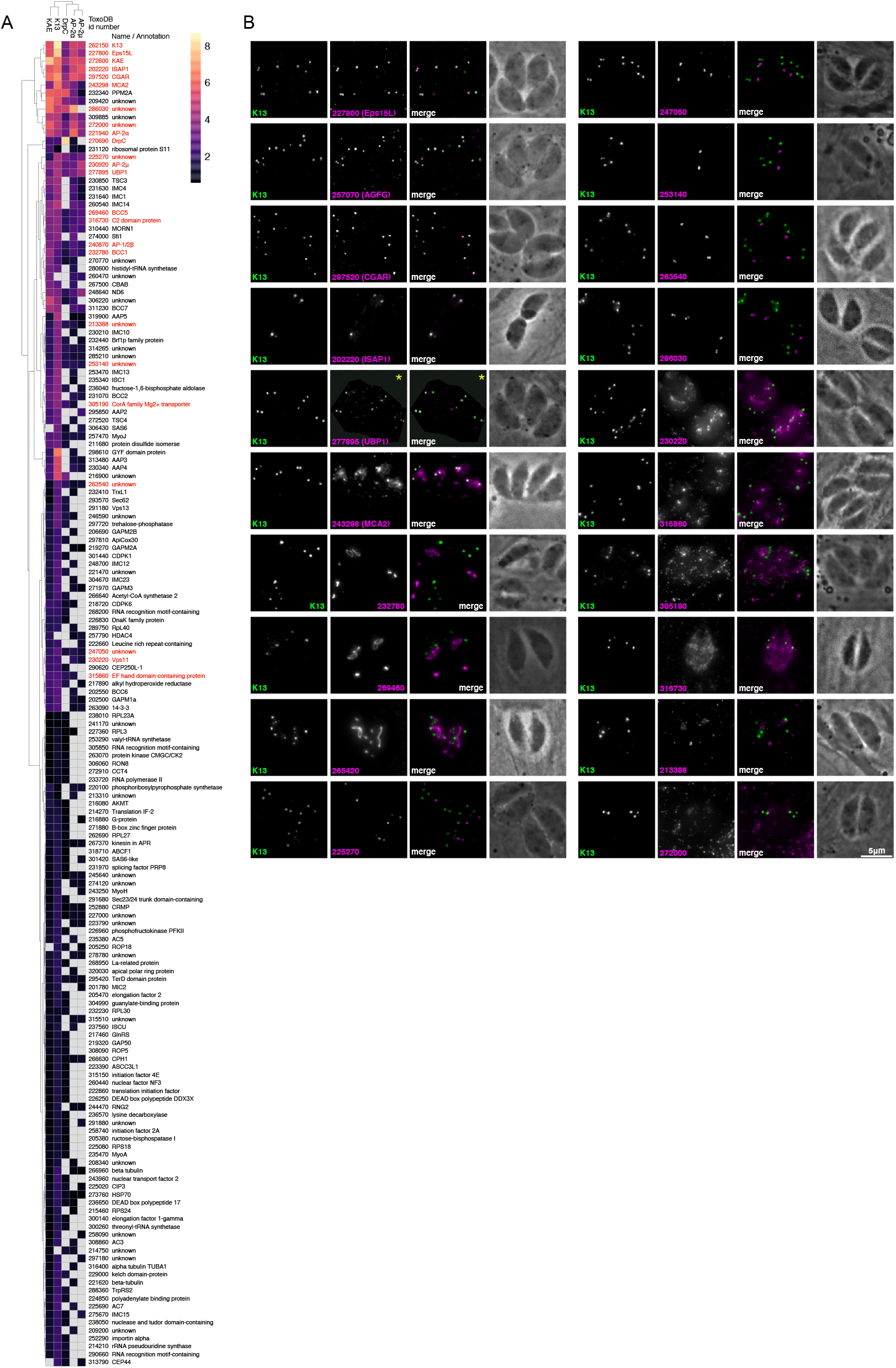
Heat map of 155 protein enrichments by the five BioID baits and microscopy validations of selected proteins. (A) Extension of plot shown in Figure 2A. The proteins that were selecting for C-terminal epitope tagging with 3x V5 tag are highlighted in red. (B) Determination of the locations of 20 proteins enriched by BioID. Six proteins show specific collocation with K13 (227800, Eps15L; 257070, AGFG; 297520, CGAR; 202220, ISAP1; 277895, UBP1), three show similarity to the basal complex (232780, 269460, 265420), four show centrosome-like positions (247050, 253140, 263540, 286030), and the remaining five protein locations are unresolved. Scale bars = 5 µm. *The area outside of the *T. gondii*-containing parasitophorous vacuole was shaded to cover a high background fluorescence, which is predominantly seen in the human HFF cells and outshines the v5 epitope signal of low abundant or less accessible proteins in *T. gondii* such as UBP1.

**Figure S4:**
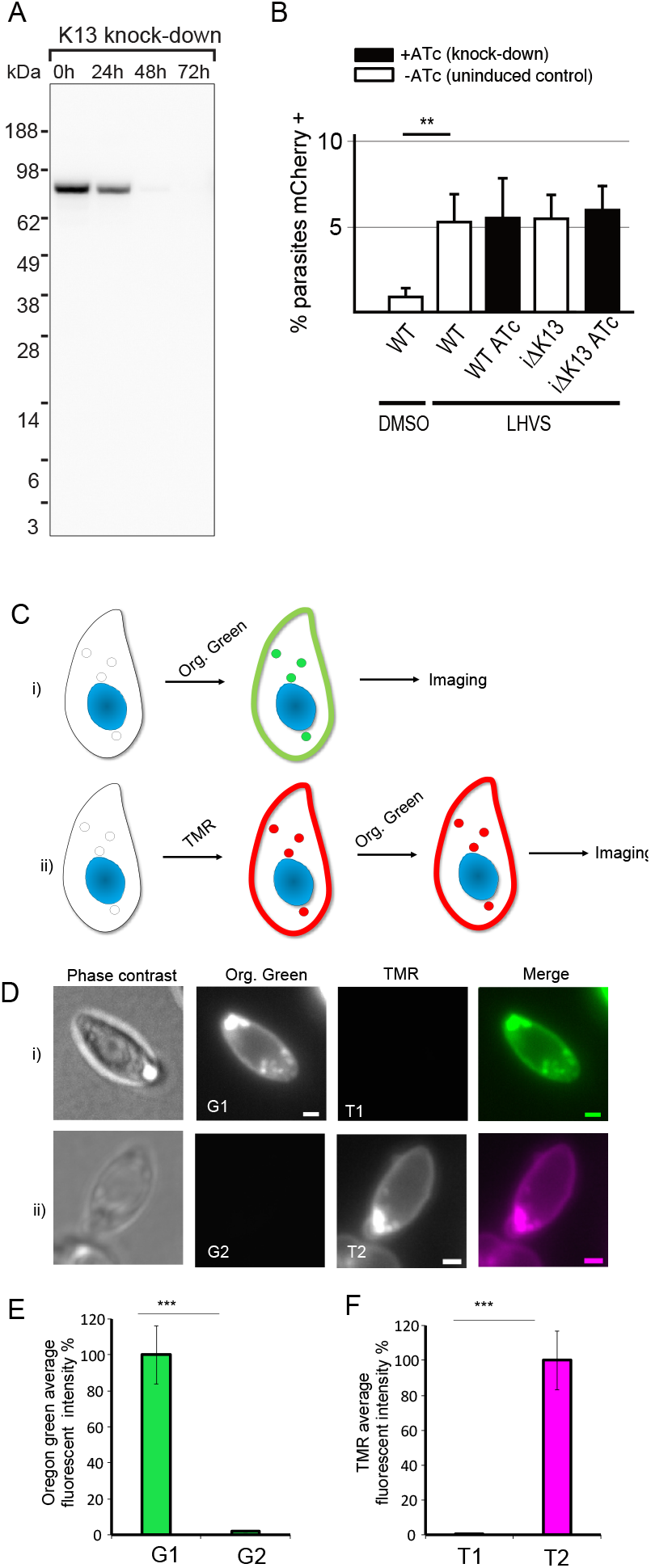
Endocytosis assays. (A) K13 depletion detected by Western blot over 72 hours of ATc-induced mRNA down-regulation. (B) Percentage of parasites with detectable host cytosol-expressed mCherry accumulation with 24 h cathepsin L inhibitor LHVS treatment or DMSO vehicle control. Parental (wild type) or iΔHA -*Tg*K13 cells were given 24 hours to accumulate mCherry following 48 hours plus or minus ATc-treatment. Histogram represent means of 4 replicates and error bars are standard deviations. ** indicates significant difference of p<0.001 by Student’s t-test. (C) Schematic of ligand binding to test for saturability of the SAG1 pools by Halo ligands. (D) i) Membrane-permeable Oregon Green labels both surface and intracellular SAG1 vesicles when used alone. ii) When membrane-permeable TMR biding is applied first, no binding of either external or internal SAG1 pools is seen by subsequent Oregon Green application. (E,F) Quantitation of parasite fluorescence for treatments according to (D). Three biological replicates were used for each analysis; P-values are indicated as 0.05<P≤1, ns; 0.01<P≤0.05, *; 0.001<P≤0.01, **; 0.0001<P≤0.001, ***. Scale bars = 1 µm.

**Figure S5:**
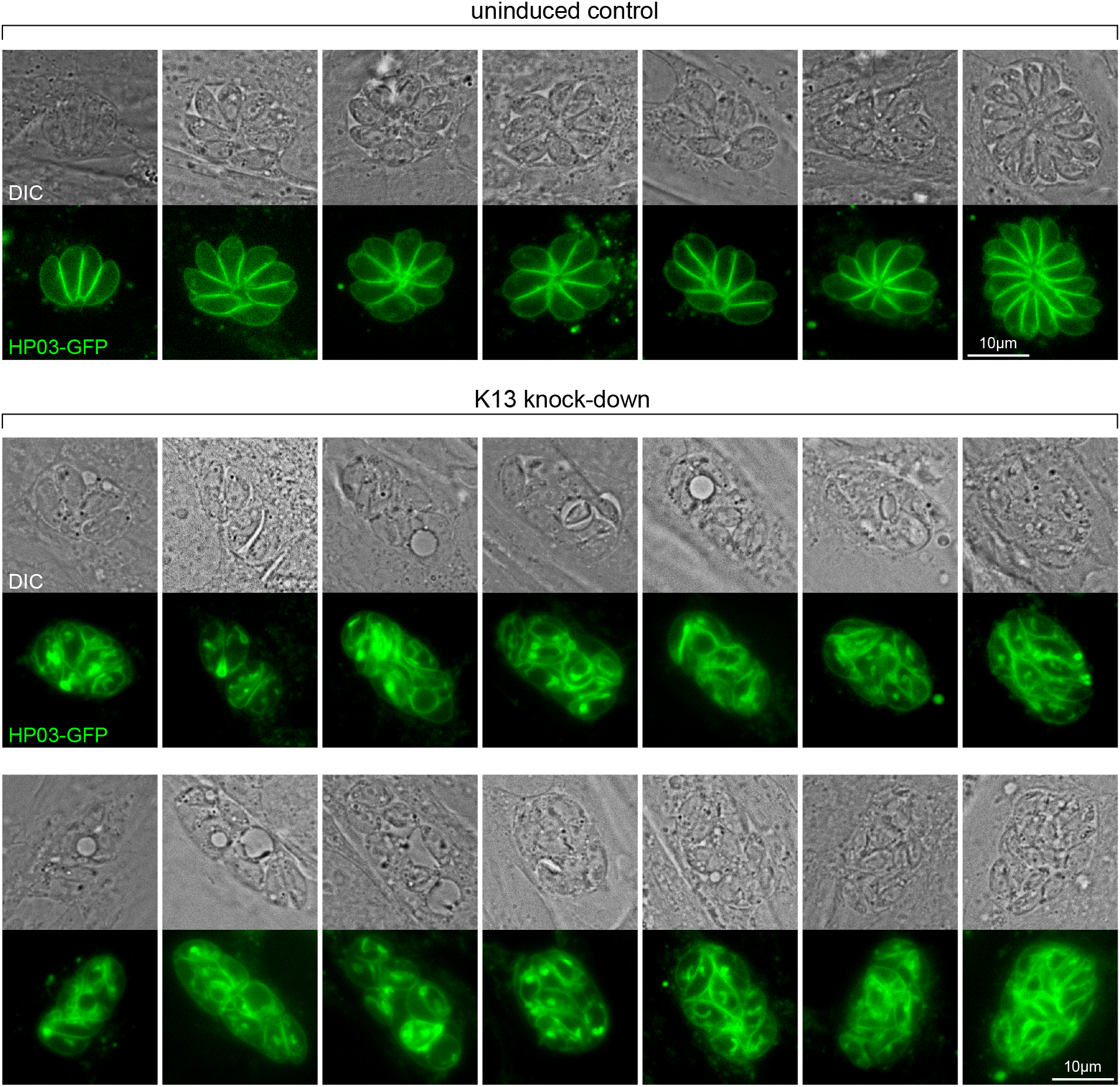
Plasma membrane and parasite rosette disruption upon K13 depletion. Live-cell microscopy by Differential Interference Contrast (DIC) and fluorescence detection of integral plasma membrane protein marker HP03-GFP shows loss of parasite organization and excess of plasma membrane. Figure an extension of Fig 6C.

**Figure S6:**
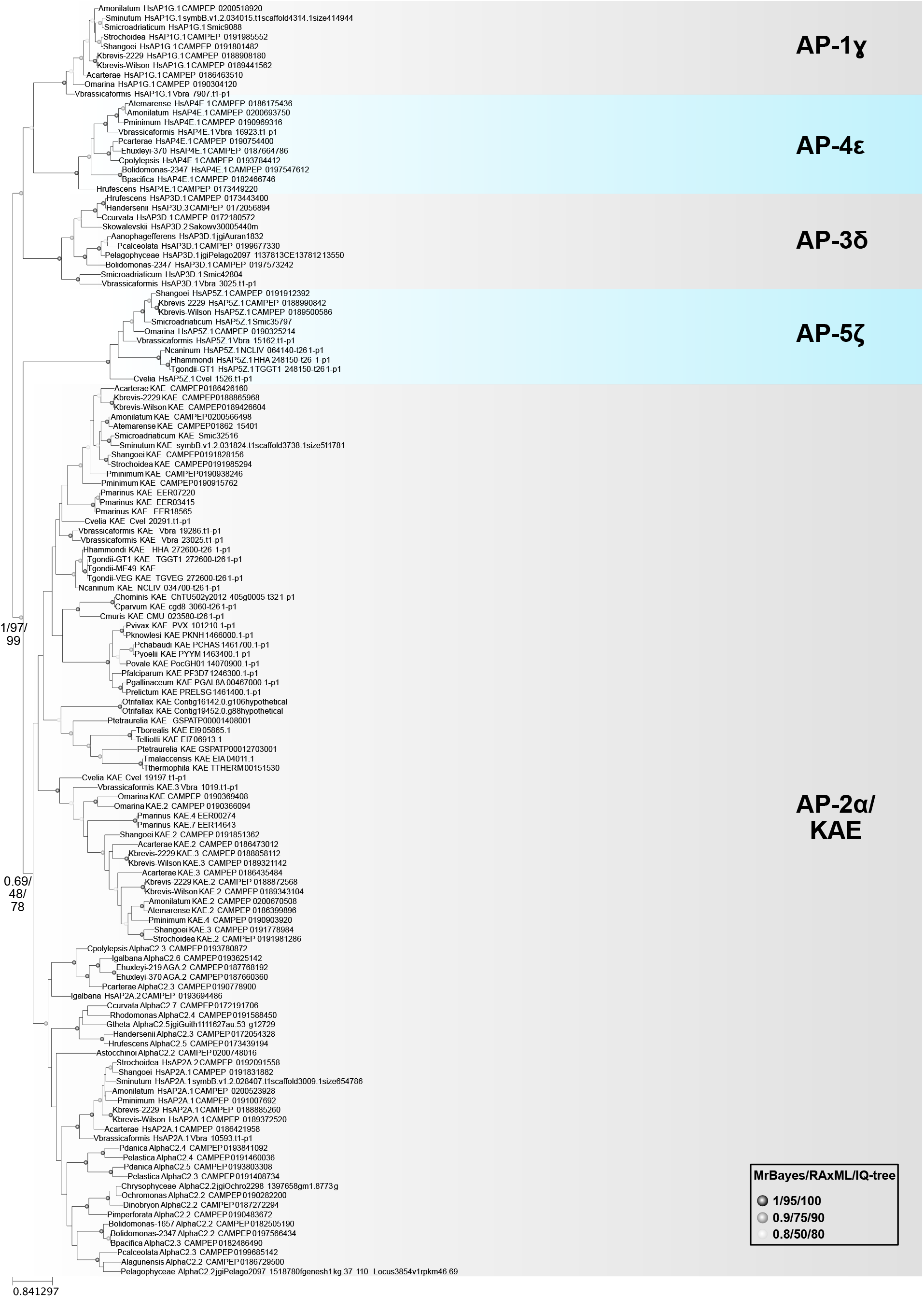
Maximum likelihood phylogeny of AP ear domains showing that KAE is derived from a duplication of AP-2α. KAE is only found in alveolate taxa and only dinoflagellates, perkinsids and chrompodellids maintain a second common KAE paralogue. Branch supports were evaluated based on MrBayes posterior probabilities and bootstrap analyses in RAxML and IQ-tree.

**Figure S7:**
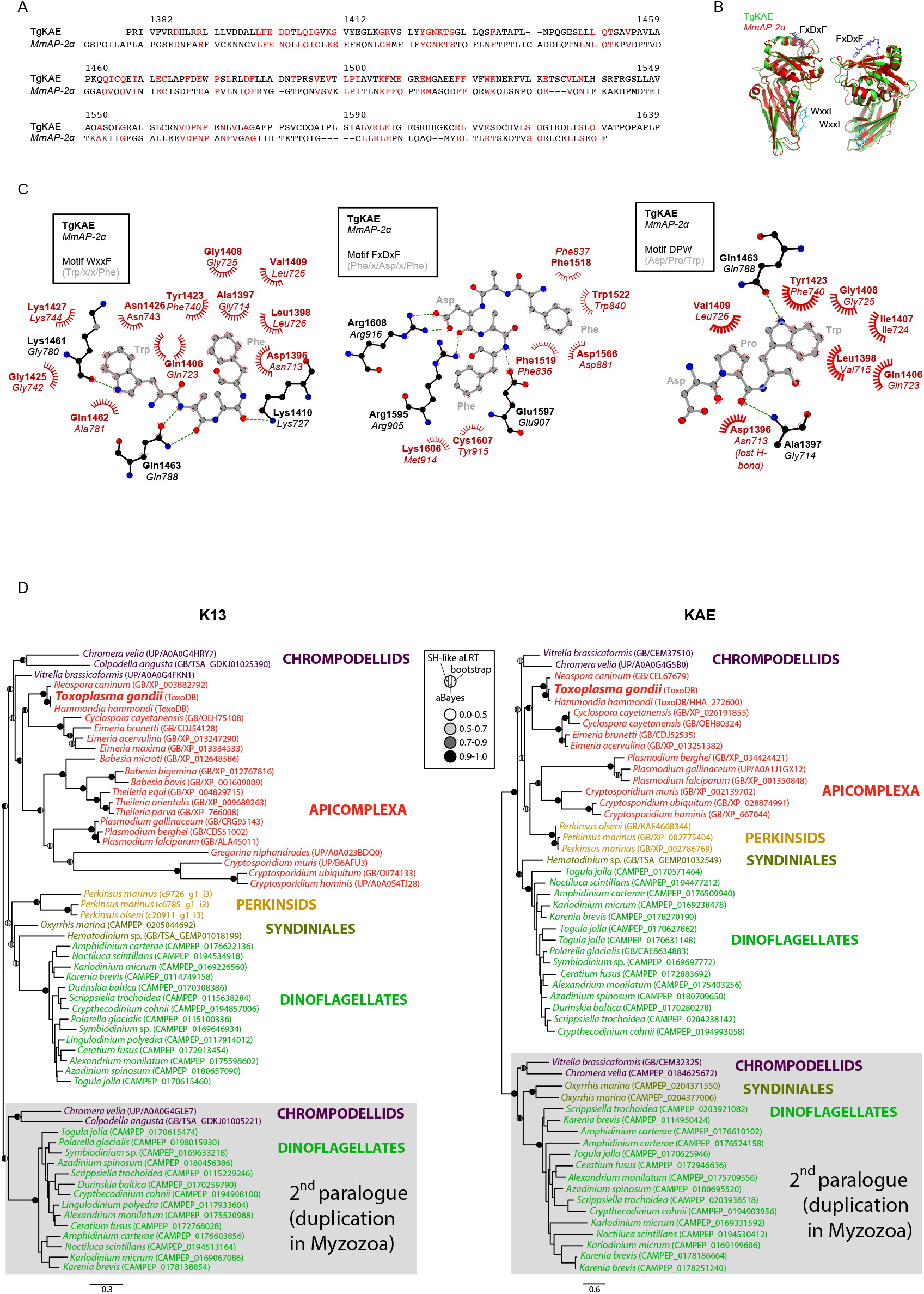
Structural and evolutionary analysis of KAE and K13. Comparison of *Tg*KAE and *Mm*AP-2α by (A) Sequence alignments of *Tg*KAE and *Mm*AP-2α ear domains (31% identity (red), 47% positive) (numbering is for *Tg*KAE), and (B) modelling based on the *Mm*AP-2α structure (PDB 1w80) indicate a strong structural conservation. The positions of FxDxF and WxxF Eps15 binding sites are shown. (C) The *Tg*KAE model indicates that the binding sites for FxDxF, DPW and WxxF motifs are likely conserved (equivalent residues in *Mm*AP-2α are shown in italics for comparison). The peptide motif and *Tg*KAE side chains are shown in ball-and-stick representation, with carbon colored grey and black respectively, oxygen in red and nitrogen in blue. Hydrogen bonds are shown as green dotted lines, while the spoked arcs represent protein residues making nonbonded contacts with the peptide motif. (D) Maximum likelihood phylogenies of K13 and KAE. The ML tree topology and branch supports (SH-like aLRT, aBayes and bootstrap, respectively) were calculated in PhyML 3.1.

